# Hypoplasticity in sensory-driven necortical circuits of Fragile X mice

**DOI:** 10.1101/2025.09.15.675519

**Authors:** Sarah Bendova, Aisling Creed, Aisha Glynne-Percy, Erin Hoeh, Owen R. Dando, Peter C. Kind, Leena E. Williams

**Affiliations:** Institute for Neuroscience and Cardiovascular Research, University of Edinburgh, EH8 9XD; Simons Initiative for the Developing Brain, University of Edinburgh, EH8 9XD

**Author notes:** Correspondence to: Leena E. Williams, Full address: The University of Edinburgh, Hugh Robson Building, 15 George Square, Edinburgh, UK EH8 9XD.

**Keywords:** Fragile X Syndrome (FXS), somatosensory cortex (S1), whisker, Long Term Potentiation (LTP), synaptic plasticity, axon initial segment (AIS)

## Abstract

Sensory experience and perceptual learning modify the receptive field properties of cortical pyramidal neurons (PNs), primarily through long-term potentiation (LTP). Repetitive sensory stimulation or rhythmic whisker stimulation (RWS), at behaviourally relevant frequencies (8 Hz), can induce long-term potentiation (LTP) of excitatory synapses in typically developing mice in the somatosensory cortex (S1).

Fragile X syndrome (FXS) is a leading single gene inherited form of intellectual disability and altered somatosensory processing is a prevalent feature. The Fragile X mouse model (*Fmr1^−/y^* mice) similarly exhibits altered somatosensory processing, including altered neuronal circuit function in S1. To assess if LTP in Layer (L) 2/3 PNs is intact upon changes in S1 circuit function we performed *ex vivo* whole-cell electrophysiology and *in vivo* two-photon calcium imaging of layer (L) 2/3 pyramidal neurons (PNs) in the S1 cortex of adult male Fragile X (*Fmr1^−/y^*) mice and typically developing (*Fmr1^+/y^*) control littermates. We found that plasticity induced by repetitive sensory stimulation was impaired in *Fmr1^−/y^* mice *ex* and *in vivo* compared to controls.

L4-evoked synaptic responses were hyperexcitable in L2/3 PNs, consistent with circuit-level disinhibition. Despite this, baseline intrinsic spiking evoked by current steps remained largely unchanged and whisker-evoked activity appeared diminished *in vivo*, which may be consistent with findings of shortened axon initial segment (AIS) length.

We also observed an increase in adaptation to repetitive stimulation in *Fmr1^−/y^* L2/3 PNs compared to controls, which may underlie failure to induce plasticity. These findings suggest that L2/3 PNs of S1 in *Fmr1^−/y^* mice are hypoplastic, potentially related to disrupted inhibitory control and rapid adaptation during repetitive sensory stimulation. Altogether, impaired LTP in the sensory cortex suggests that the receptive field properties of cortical PNs are less flexible and dynamic, which may contribute to altered somatosensory processing and reactivity in FXS.

## Introduction

Sensory experience and perceptual learning change the receptive field properties of pyramidal neurons (PNs) in the cortex ^1–19^. In large part, Long Term Potentiation (LTP) of synapses mechanistically underlies these changes in receptive field dynamics and neuronal circuit plasticity^1,20^.

Rodents explore their environment by moving their whiskers over surfaces in rhythmic sweeps (5-15 Hz) and spiking in the somatosensory cortex (S1) is modulated in synchrony with whisking ^21–24^. Cortical representation in layer (L) 2/3 PNs of S1 is dynamic and can be modulated over an extended period via LTP-driven by rhythmic whisker stimulation (RWS), at rates for which rodents take in sensory information (8Hz) ^11,19,25^. When we dissected the neuronal circuitry underlying L2/3 PN LTP we found that this was driven by the paired activation of incoming sensory information through cortical L4 and higher-order inputs from the Posterior Medial (POm) Complex of the Thalamus, which may relay contextual signals ^25–31^. This type of LTP-dependent plasticity may serve as a mechanism to allow contextual information to shape incoming sensory information, consequently, impaired plasticity expression in S1 could disrupt tactile perception and perceptual learning.

Fragile X Syndrome (FXS) is a leading inherited form of intellectual disability due to loss of Fragile X Messenger Ribonucleoprotein (FMRP) and a core feature is altered somatosensory processing ^32–42^. The FXS mouse model (*Fmr1^−/y^* mice) is well characterized and similarly exhibits altered somatosensory processing, including changes in neuronal circuit function in S1^43–48^. To assess whether LTP and the modulation of cortical representation in S1 is impacted by altered S1 processing we employed whole-cell patch clamp electrophysiology in thalamocortical slices and monitored whisker-evoked calcium (Ca^2+^) signals in L2/3 neurons using two-photon laser scanning microscopy (2P) in S1 of *Fmr1^−/y^*mice and littermate controls.

We found both *ex* and *in vivo* plasticity in L2/3 neurons of S1 is impaired in *Fmr1^−/y^* mice. When we examined subpopulations of the notably heterogeneous L2/3 population we observed a hypoplasticity specific to persistent cells, which exhibited whisker-evoked events across the experiment. In *Fmr1^−/y^* brain slices we observed enhanced L4 to L2/3 PN subthreshold hyperexcitability that may be attributable to a lack of inhibition. Intrinsic excitability was unchanged in *Fmr1^−/y^* mice and there was a modest decrease in the number of cells activated by whisker stimulation. We observed a shortening of AIS length in L2/3 of *Fmr1^−/y^*mice ^49–54^. We also observed an increase in adaptation to repetitive stimulation in *Fmr1^−/y^* L2/3 PN compared to controls, which may underlie the failure to induce plasticity.

## Materials and methods

### Mice

*Fmr1-KO* mice (B6.129-Fmr1tm1Cgr, RRID: MGI_MGI:3815018) were obtained from the Contactor laboratory (Northwestern University, Chicago IL, USA)^43^. Mice were maintained on a C57Bl6/J background. Hemizygous male *Fmr1-KO* (*Fmr1^−/y^*) and control littermates (*Fmr1^+/y^*) were group housed on a 12-hour light/dark cycle. All procedures were carried out according to UK Home Office and NIH IACUC guidelines for animal welfare.

### Brain slice preparation

6–8-week-old mice were anesthetized with isoflurane and decapitated. A vibrating microtome was used to prepare 350-mm-thick thalamocortical slices according to a method for adult and aging animals ^55,56^. Slicing was performed in cold NMDG artificial cerebrospinal fluid solution (aCSF, 300-305 mOSm, pH7.3), containing (in mM): 92 NMDG, 2.5 KCl, 1.25 NaH_2_PO4*H_2_O, 30 NaHCO_3_, 20 HEPES, 25 glucose, 2 thiourea, 5 Na-ascorbate, 3 Na-pyruvate, 0.5 CaCl_2_*2H20, 10 MgSO_4_*7H_2_O ^55,56^. Slices were transferred to NMDG aCSF solution at 35°C for 20 min and immersed in room temperature HEPES aCSF solution (300-305mOSm, pH 7.3, in mM): 92 NaCl, 2.5 KCl, 1.25 NaH_2_PO_4_*H_2_O, 30 NaHCO_3_, 20 HEPES, 25 glucose, 2 thiourea, 5 Na-ascorbate, 3 Na-pyruvate, 2 CaCl_2_*2H_2_O, and MgSO_4_*7H_2_O. 95%, O_2_ +5%CO_2_ was bubbled though all solutions.

### Electrophysiology

Whole-cell patch-clamp recordings were obtained from L2/3 PNs. Recordings were performed in freshly prepared aCSF, bubbled with 95% O_2_ +5%CO_2_, at an osmolarity of 300-305 mOsm, containing (in mM): 119 NaCl, 24 NaHCO_3_, 2.5 KCl, 1.25 NaH_2_PO_4_*H_2_O, 2 MgSO_4_*7H_2_O, 2 CaCl_2_*2H_2_O, and 12.5 Glucose. All recordings were performed at 34 ± 1 °C. Infrared differential inference contrast (IR-DIC) video microscopy, using a digital camera (Qimaging) mounted on an upright microscope (Olympus BX51WI) and a 40 × (0.8NA) water immersion objective. Recordings were obtained with a Multiclamp 700B (Molecular Devices) amplifier, signals were Bessel filtered (1 kHz) online and digitized at 20 kHz (Digidata1440, Molecular Devices) coupled to the Clampex software (pCLAMP™ Software, Molecular Devices). Recording pipettes were pulled from borosilicate glass capillaries (1.7 mm outer/1 mm inner diameter, Harvard Apparatus, UK) on a horizontal electrode puller (P-97, Sutter Instruments, CA, USA).

Patch pipettes (4-9 MΩ) were filled with 290-305 mOsm internal solution containing (in mM): 135 K-gluconate, 5 KCl, 10 Phosphocreatine, 4 Mg ATP, 0.3 NaGTP, 2.68 (0.1%) Biocytin, and 10mM HEPES. After break-in the cell equilibrated for 5 minutes. Membrane capacitance (Cm) and series resistance (Rs) was monitored throughout, and cells were discarded when Rs changed >20%. A series of current injection (40 pA increments, 500 ms) steps were applied to measure intrinsic properties and spike patterns to identify PNs specifically.

All electrophysiological recordings and analysis were performed using pClamp 11 software (Molecular Devices). All mEPSCs, sEPSCs, EPSCs, and PSPs were detected and measured using Event Analysis tools in pClamp 11 software. Amplitude and frequency measurements are averages from all events occurring within 120 sec of continuous recording.

#### LTP protocol

For plasticity experiments, L4-electrical stimulation (L4-ES) was tuned to yield a 5mV Post Synaptic Potential (PSP) baseline (pre) response. The LTP protocol consisted of a 5-min baseline period of L4-ES (0.1Hz), followed by a 1-min period of rhythmic electrical stimulation (RES; 8Hz), and a 30-min plasticity readout with the same baseline stimulation. We compared average baseline PSP amplitudes (pre L4-RES, 0-5 min; 30 stimulations) to those over the final 5 mins of recording (post L4-RES, 25-30 min; 30 stimulations). The level of LTP was calculated per cell and mouse. Synaptic responses were monitored before, during, and after the L4-RES.

#### Evoked, minimal stimulation, & paired-pulse ratio

L2/3 PN PSPs and Excitatory Postsynaptic Currents (EPSCs) were evoked by L4-electrical stimulation (L4-ES, 0.2 ms) with a bipolar stimulating electrode (matrix tungsten electrode, FHC) placed in L4. For the PSP and EPSC input/output (I/O) curve, responses were recorded across increasing L4-ES intensities (50, 100-900 µA). To block inhibition 10 µM bicuculline (to block GABAA receptors) was bath applied. For minimal stimulation experiments, once a reliable EPSC was identified the stimulus intensity (50-1000 μA) was reduced until the synaptic response fail in ∼half of the trials, allowing for the putative stimulation of single or a very few presynaptic fibres ^57^. For paired pulse measurements two L4-electrical stimuli were given in rapid succession across four interstimulus intervals (ISIs, 10, 20, 50 100 ms) and the difference in response sizes to the two stimuli were calculated as a ratio (1^st^/2^nd^ EPSC).

#### mEPSCs and sEPSCs

mEPSCs recordings were performed voltage clamped at −70mV in bath solution containing 10 µM bicuculline and 1 μM TTX (to block presynaptic action potentials) with 5 μM cyanquixaline (CNQX) as a control.

### Immunohistochemistry

Mice (8-10 weeks) were terminally anaesthetised using intraperitoneal injection of sodium pentobarbital (200 mg/ml; Dolethal, Vetoquinol) and were transcardially perfused with PBS followed by 4% PFA (Sigma-Aldrich) in PBS.

#### Immunohistochemical quantification of synaptic proteins

Whole brains were fixed in 4% PFA for 24 hours at 4°C and cryoprotected by submersion in 30% sucrose in PBS solution for 48 hours. Coronal brain sections (40 μm thickness) were obtained using a freeze sliding microtome (HM 450, Thermo Scientific) and stored in PBS. Coronal sections were washed three times in PBS, blocked for 1 hour in 0.1 M PBS containing 0.025% Triton X-100 and 5% bovine serum albumin (BSA), and incubated at 4°C for 18-24 hours in blocking solution with primary antibodies; rabbit anti-glutamate receptor 1 of AMPA subtype phospho T840 (T840-P GluA1, 1:1000; Abcam #12108, RRID: AB_2113448) and mouse anti-postsynaptic density protein-95 (PSD95, 1:500; Invitrogen #6G6-1C9, RRID: AB_325399). The sections were washed four times in PBS with 0.025% Triton X-100 and incubated at 4°C for 2 hours in PBS with 5% BSA and appropriately fluorescently conjugated secondary antibodies (1:400; anti-rabbit and mouse polyclonal IgG with AlexaFluor-488 and - 647 Thermo Fisher, A-11008, RRID: AB_143165; or 11001, RRID: AB_2534069; and anti-rabbit or mouse AlexaFlour568, 1:400; Thermo Fisher, A-11004, RRID: AB_2534072; or 11011, RRID: AB_143157). Secondary fluorophores were switched in half of the animals as a control. Sections were washed three times in PBS, mounted on glass slides, covered with Vectashield antifade mounting medium (Vector Laboratories), and secured with a glass coverslip sealed by a nail varnish. Negative controls were stained with fluorescent secondary and no primary antibodies.

Confocal microscopy (Nikon A1R FLIM with 60x, 1.4 NA, oil-immersive objective lens; Nikon) was used to generate a 540x magnification z-stack image with z-interval of 0.175 μm across 20 μm, from the L2/3, 100-300 µm from pia, and three images per animal (61.4 x 61.4 μm, 1024 x 1024 pixels, pixel size: 0.06).

Colocalized overlapping T840-P GluA1 and PSD-95 puncta were identified in this region and puncta edges were manually delineated using the freehand tool, across the z stack. The size and density of the puncta were calculated using ImageJ by measuring the area dimensions and mean fluorescence intensity. Measurements were averaged per each section and across the three images, generating an average T840-P GluA1 and PSD-95 synaptic puncta size and density per animal.

#### Immunohistochemical quantification of AIS length

The brain was removed and post-fixed in 4% PFA for 2 h (to optimize Ankryin-G staining) at room temperature, before transfer into PBS and stored at 4°C. Brains were cryoprotected in 30% sucrose in PBS, for 48 h prior to slicing ^58^. Coronal sections (100 µm) were sliced using a cryostat (Leica CM3050 S) allowing brains to freeze in Optimal Cutting Temperature (OCT) 20 mins before slicing. Slices were stored in PBS at 4°C. Free floating slices were washed with PBS for 10 minutes, 4 times, on a shaking plate at room temperature. Slices were then blocked with 10% normal goat serum (NGS), 0.3% Triton X-100, 0.05% NaN3 in PBS for 1 h at room temperature on a shaking plate. Primary antibodies, anti-Ankyrin-G (mouse monoclonal, 1:500; Neuromab, N106/36; RRID: AB_1067303), an AIS marker, and anti-NeuN (rabbit polyclonal, 1:500; Millipore EMD, MAB377; RRID: AB_10807945) were added along with 5% NGS, 0.3% Triton X-100 and 0.05% NaN_3_ diluted in PBS, for 24 hours at 4°C ^58^. Slices were then thoroughly washed. Secondary antibodies (anti-rabbit or mouse AlexaFluor488, 1:400) were applied in wells containing 3% NGS, 0.1% Triton X-100 and 0.05% NaN3 diluted in PBS, for 24 h at 4°C. Slices were washed, mounted on glass slides, and Vectashield mounting medium was applied.

Z-stacks were collected with (Leica TCS SP8) invert scanning-confocal microscope with a 63x (HC PL APO 63x CS2, N.A. 1.4, oil-immersion) objective. The S1 images were taken for L2/3 100-300 μm or L4 300-500 μm from the pia. Z-stacks (0.3μm steps, 1024×1024 pixels, scan speed 600 Hz) of the middle 20 μm of the slice were imaged. Two Z-stacks images from two different slides were taken per mouse for analysis.

All images were analysed in the FIJI package of ImageJ. AIS lengths were traced manually using the segmented line tool in FIJI and the anti-Ankyrin-G (AlexaFlour568/AlexaFlour488) labelling, from the most distal tip to the most proximal end of the AIS ^54^. AISs on the edge of Z-stacks or end of plane were excluded.

### Surgery and virus injections

Stereotaxic injections of adeno-associated viral (AAV) vectors were carried out on 7-week-old male mice. A mix of O_2_ and 4% isoflurane at _∼_0.4 L min^−1^ was used for anesthesia. AAV2-CAG-GCaMP6s-WPRE-SV40 (UPenn Vector Core, RRID: Addgene_100844, 100 nl), or AAV1-hSynmRuby2-GSG-P2A-GCaMP6s-WPRE-pA (Addgene, RRID: Addgene_ 50942; 100 nl) was delivered to L2/3 of the right barrel cortex at the approximate location of the C2 barrel column (1.4 mm posterior, 3.5 mm lateral from the bregma, 300 mm below the pia; Rose et al., 2016). A 3-mm-diameter cranial window was implanted, as described previously ^59^.

### RWS protocol

Eyes were covered with a thin layer of transparent eye ointment (Viscotears gel) and head-fixed to the two-photon set up. Body temperature was kept constant at 37 via a temperature control system (Harvard Apparatus). Mice were anesthetized with intraperitoneal injection of MM solution, consisting of 0.2 mg kg^−1^ medetomidine (Domitor, Orion Pharma), and 5 mg kg^−1^ midazolam (Dormicum, Roche) diluted in sterile 0.9% NaCl. For all experiments pre- and post-RWS, all whiskers were deflected back and forth (five deflections lasting each 45 ms @ 20 Hz) for 10 min at a frequency of 0.1 Hz. Whiskers were deflected with a piezoelectric ceramic element attached to a glass pipette with a 3D printed grid 4 mm away from the skin. The voltage applied to the ceramic was set to evoke a whisker displacement of 0.6 mm with a ramp of 7–8 ms.

### *In vivo* two-photon imaging

Imaging was performed using a custom-built resonant scanning two-photon microscope as described previously. In brief, the set up was equipped with a Ti:Sapphire excitation laser (Charmeleon Vision-S, Coherent, CA) at 920 nm and GaAsP photomultiplier tubes (Scientifica). Images were acquired at 20 Hz with a custom-programmed LabView based software (v8.2; National Instruments, UK) using a Nikon 40x water-immersion objective (Nikon; CF175 Apo 25XC W; 1.1 NA). Time-series images of one focal plane (field of view; FOV) per mouse were acquired at cortical depths between 100 and 300 μm from pia.

### Data analysis

Motion correction was conducted using Non-Rigid Motion Correction (NoRMCorre) algorithm ^60^. This was done per recording and across all the recordings. For outlining ROIs, the motion-corrected recordings were each averaged to one frame per each recording and opened in ImageJ to manually define individual neurons in the GCaMP6s channel with the aid of the mRuby2 channel, an activity independent marker, using the freehand draw tool. For extraction of D (Delta)F/F_0_ we used FISSA ^61^. To remove neuropil contamination, we used Fast Image Signal Separation Analysis (*FISSA*149) which uses non-negative matrix factorization (NMF) to demix spatiotemporal overlapping signal sources through blind source separation. The relative decontaminated fluorescence intensity of each ROI was calculated as ΔF/F_0_ (or as (F−F_0_)/F_0_), where F_0_ is the 5^th^ percentile of the 1 Hz low pass filtered signal across the entire imaging session, and ΔF is the subtraction of F_0_ from the decontaminated fluorescence intensity of an ROI in a frame. Python toolboxes (*SIMA* and *FISSA*) were run with WinPython 2.7.10.3. Further analysis was performed with custom-written scripts in MATLAB (MathWorks, MA). Firstly, for each ROI during the recordings from the pre and post-RWS period, the extracted ΔF/F_0_ signals were detected as Ca^2+^ events if the increase in ΔF/F_0_ signal was 2.5 times its standard deviation with a 0.25 sec rise time threshold. The Ca^2+^ events were then aligned with the whisker stimulation triggers and considered a whisker-evoked Ca^2+^ event if the rise time was within 1 sec from the stimulation onset.

For each neuron in the pre- and post-RWS period the signal probability (PE), the mean amplitude (*A_E_*), and the response strength (RS_E_) of the whisker-evoked Ca^2+^ events were calculated as follows with being the number of whisker-evoked Ca^2+^ events, the overall number of stimulations, and *A_E_* the amplitude of a whisker-evoked Ca^2+^ event:

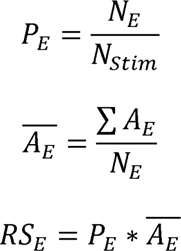

Mean fluorescence intensity during RWS (DF/F0) was calculated, and the Ca^2+^ fluorescence intensity was integrated over the baseline (20 s before start of RWS) and during the RWS period (20 s after the start of RWS)19.

### Experimental design and statistical analysis

All statistics were performed as reported in the text and figure legends, and graphs were created using Prism 9 or 10 (GraphPad Software). For all figures, significance levels were denoted as *p<0.05, **p<0.01, ***p<0.001, and****p<0.0001, and asterisks were reported per mouse in the figures. All data are reported as mean±standard error of the mean (SEM) per mouse reported in the figure legends. In electrophysiology and AIS figures mice averages are represented in bar graph and pairwise comparisons. For 2-photon figures cell averages are represented in bar graph and pairwise comparisons, whereas mice (in figures) averages are represented as adjacent circles±SEM. For all experiments, n equals the number of cells and N equals number of mice. All data were assessed for normality, if the data passed normality a Paired or Unpaired t test was performed. If not, a Wilcoxon matched-pairs signed rank test or Mann-Whitney test was performed. For comparison of distributions a Kolmogrov-Smirnov (KS) test was performed. For comparisons of cell population proportions, a binomial or Chi square test was performed. All tests were performed two sided.

For data with multiple measurements per animal, generalized linear mixed modelling (GLMM) was performed to account for random effects in RStudio v2024.12.1 and were reported in the text and in Supplemental Values and Statistics. Any discrepancy between traditional statistical tests per mouse methods and the GLMM were highlighted. Depending on the distribution, either linear mixed models (LMMs) or GLMMs were fit using the R package lme4 v1.1.37. In the event of multinomial count data with random effects, a GLMM was fit using the R package mclogit v0.9.6. For comparisons between genotypes, such as in Figure 4E, models include an animal random effect and a genotype fixed effect. In other cases, such as comparing response strength in Figures 5D and 5E, a per genotype effect size was of interest and those models were fit with an animal random effect and an intercept term. P-values for testing single parameter significance are given by Wald Z-tests. For testing of effects, p-values are given by likelihood ratio tests comparing a model containing the variable of interest and the null model without that variable. On occasion, the data was log-transformed to meet model assumptions of normality and an LMM was fit to the data. Consequently, regression coefficients are provided on log scale.

All analysis was performed blinded to the genotype. No data were excluded. For the assessment of LTP *ex vivo*, glutamate receptors, Ca^2+^ imaging, and AIS measurements *a priori* power analyses were performed to estimate sample size using G*Power (version 3.1.9.7.) based on published data with a sample size statistical power of >83% at 0.05 significance level ^19,27,54,62^.

## Results

### Rhythmic L4 -evoked plasticity is impaired in *Fmr1^−/y^ brain slices*

Previous work indicates that rhythmic activation of L2/3 synapses can induce LTP ^2,11,19,25,27,63–65^. To test if L2/3 PN LTP is intact in *Fmr1^−/y^* mice we tested the effect of a brief (1 min) period of rhythmic (8 Hz) electrical stimulation of L4 (L4-RES) in brain slices from *Fmr1^−/y^* and *Fmr1^+/y^* littermate controls and measured L4-ES PSP amplitudes pre and post (Fig. 1 A). We found that L4-RES significantly increased mean post control *Fmr1^+/y^*L2/3 PN PSP amplitudes (*P*=0.008, Fig. 1B, D). In contrast, mean post *Fmr1^−/y^*L2/3 PN PSP amplitudes remained unchanged (*P*=0.6, ANOVA *P*=0.008, Fig. 1 C, D). We found no difference in L2/3 PN PSP size pre between genotypes (Supplementary Fig.1). We found that significantly less L4-electrical stimulation (ES) current was necessary to evoke similar PSP sizes in *Fmr1^−/y^* L2/3 PN compared to littermates, but there was no correlation between L4-ES current and the size of the LTP (Supplementary Fig. 1). Together this demonstrates that LTP activated by L4-RES in L2/3 PNs is impaired in brain slices from *Fmr1^−/y^* mice compared to controls.

**Figure 1.**
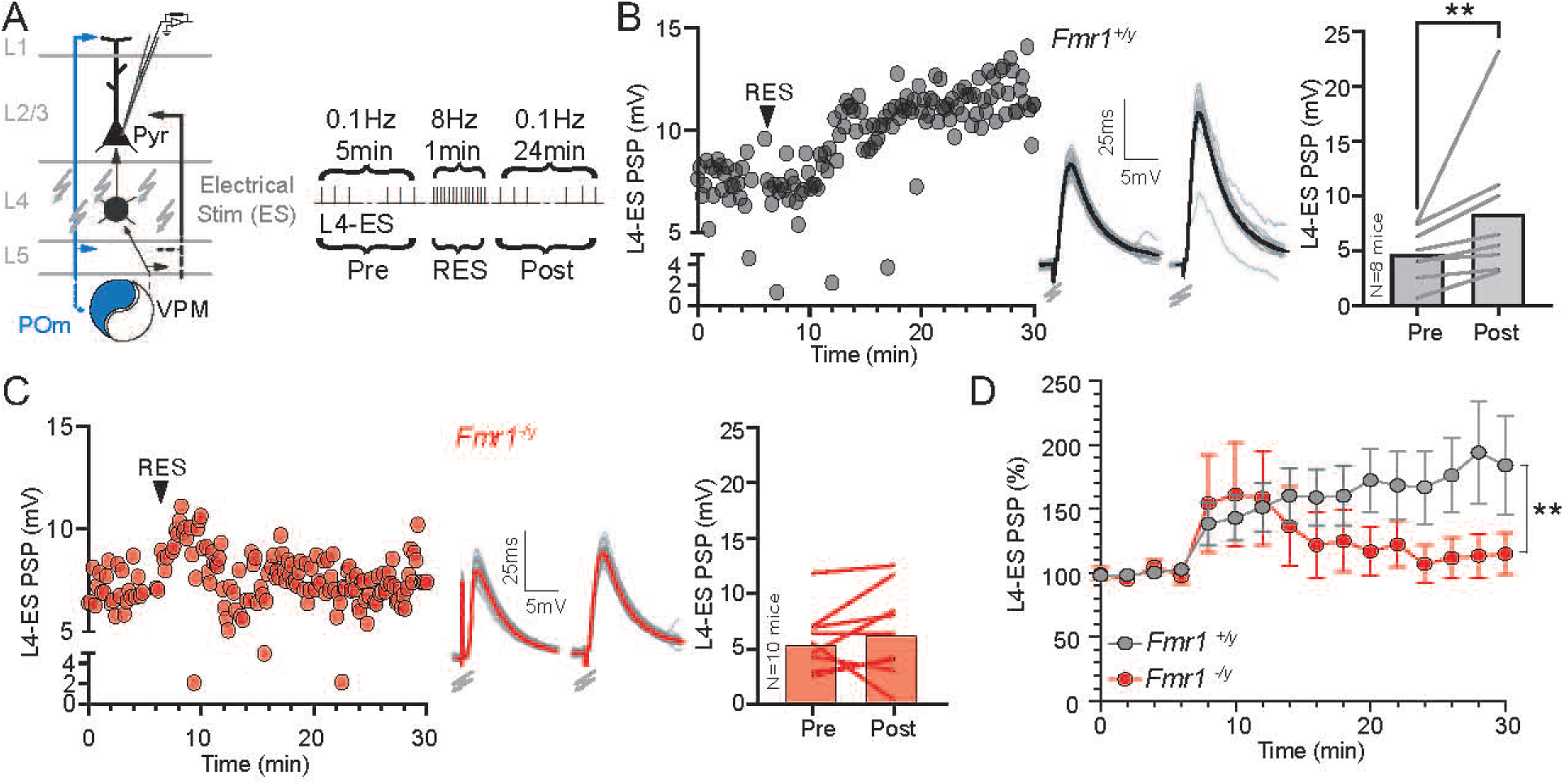
Rhythmic L4 -evoked plasticity is impaired in in *Fmr1^−/Y^* mice. **(A)** Left, schematic of the somatosensory thalamocortical (POm and VPM) projections and their relation to intracortical inputs in the BC. Recordings are made in L2/3 PNs while electrically stimulating L4 (L4-ES). Right, experimental protocol: L4-ES at 0.1 Hz (pre-RES; 5 min), followed by rhythmic electrical stimulation (RES) at 8 Hz (1 min), followed by L4-ES at 0.1 Hz (post-RES; 24 min). **(B)** Left, L4-ES PSP amplitudes in an example *Fmr1^+/Y^* cell. Middle, representative L4-ES PSPs pre- versus post-RES are shown. Right, mouse averages (bars) and pairwise connections (lines) L4-ES PSP amplitudes pre- versus post-RES (*N*=8, mean +/− SEM, pre=4.72+/−0.85 mV, post=8.41+/−2.35, *P*=0.008). **(C)** Left, L4-ES PSP amplitudes in an example *Fmr1^−/Y^* cell. Middle, representative L4-ES PSPs pre- versus post-RES are shown. Right, the population (bars) and cell (lines) mean L4-ES PSP amplitudes pre- versus post-RES (*N*=10, pre=5.47+/−0.89 mV, post=6.28+/−1.23, *P*=0.6). **(D)** Normalized L4-ES PSP amplitudes in L2/3 PNs after RES in *Fmr1^+/Y^* (*N*=8) & *Fmr1^−/y^* (*N*=10) brain slices (Two-way ANOVA, F (15, 240) = 2.164, time x genotype *P*=0.008).

### Hyperexcitable L4 to L2/3 PN inputs in *Fmr1^−/y^* mice

To examine inputs to L2/3 PNs and probe causality of impaired LTP we recorded in current clamp L2/3 PN PSP responses to single L4-ES at various stimulation currents (50-900µA) to generate an input/output (I/O) curve. We found a significant increase in PSP size across the I/O curve and half-max amplitude in *Fmr1^−/y^* compared to control L2/3 PNs (ANOVA *P*=0.02, *P*=0.009, Fig. 2A). In separate voltage clamp experiments, we again found a significant increase in L2/3 PN EPSCs and half-max for *Fmr1^−/y^* compared to control L2/3 PNs (ANOVA *P*<0.0001, *P*=0.0006, Fig. 2B).

**Figure 2.**
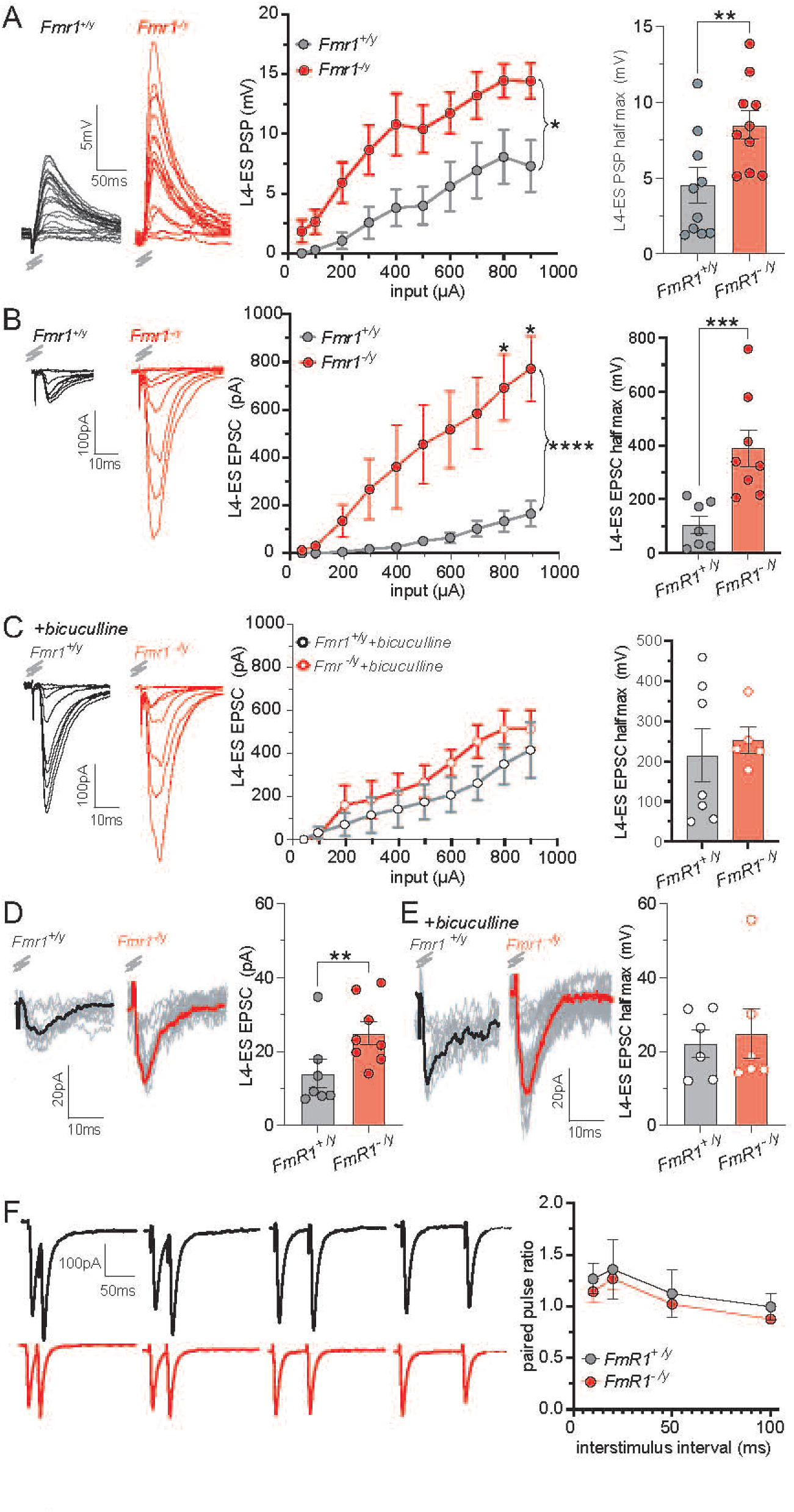
Hyperexcitable L4 to L2/3 PN inputs in *Fmr1^−/y^* mice. **(A)** Left, representative L4-ES PSPs and middle, amplitudes across various L4-Electrical stimulation (ES) inputs (50, 100-900 µA) for *Fmr1^+/y^* & *Fmr1^−/y^* L2/3 PNs (Two-way ANOVA, genotype, F(1, 17)= 6.6, *P*=0.02). Right, L4-ES PSP half-max amplitude for *Fmr1^+/y^* (N=10 mice, 4.31+/− 1.10) & *Fmr1^−/y^* (N=10 mice, 8.48 +/− 0.94, *P*=0.009) L2/3 PNs. **(B)** Representative L4-ES EPSCs for *Fmr1^+/y^* and *Fmr1^−/y^* L2/3 PNs. Middle, EPSCs across various L4-ES inputs for *Fmr1^+/y^* & *Fmr1^−/y^* L2/3 PNs (Two-way ANOVA, time x genotype, F(9, 126)= 7.9, p<0.0001; Šídák’s multiple comparison, only significant data reported, 800µA p=0.04, & 900µA p=0.02). Right, L4-ES EPSC half-max amplitude for *Fmr1^+/y^* (N=8 mice, 104.5+/− 32.3) & *Fmr1^−/y^* (N=8 mice, 388.6 +/− 67.8, *P*=0.0006) L2/3 PNs. **(C)** Representative L4-ES EPSCs from *Fmr1^+/y^* and *Fmr1^−/y^* L2/3 PNs with bath applied bicuculline. Middle, EPSCs across various L4-ES inputs for *Fmr1^+/y^*& *Fmr1^−/y^* L2/3 PNs (Two-way ANOVA, time x genotype, F(9, 90)=0.7, *P*=0.7). Right, L4-ES half-max amplitude for *Fmr1^+/y^* (N=7 mice, 214.4+/− 66.2) & *Fmr1^−/y^* (N=5 mice, 252.5 +/− 32.5, *P*=0.7) L2/3 PNs. **(D & E)** Left, representative minimal L4-ES EPSCs from *Fmr1^+/y^* & *Fmr1^−/y^* L2/3 PNs without (D) & with bicuculline (E). Right, mean minimal L4-ES EPSC amplitudes from *Fmr1^+/y^* & *Fmr1^−/y^* L2/3 PNs without (D, *Fmr1^+/y^*=13.2+/−3.4, N=8; *Fmr1^−/y^* =25.1+/−3.1, N=8, p=0.007) & with (E, *Fmr1^+/y^*=20.3+/−3.6, N=7; *Fmr1^−/y^* =24.9+/−6.6, N=6, *P*=0.6) bicuculline. **(F)** Left, Representative paired pulse L4-ES EPSCs for *Fmr1^+/y^*and *Fmr1^−/y^* L2/3 PNs across four interstimulus intervals (ISI, 10, 20, 50, 100ms). Right, ISI versus paired pulse ratio (post/pre %, Two-way ANOVA, ISI x genotype, *P*=1.0).

Next, to test if increased *Fmr1^−/y^* L4 to L2/3 PN synaptic responses are causally linked to inhibition we bath applied bicuculline (10 μM), the GABAA receptor antagonist, and examined L2/3 PN EPSCs, which eliminated genotypic differences (ANOVA *P*=0.7, Fig. 2 C). For control L2/3 PNs the addition of bicuculline resulted in a significant increase in EPSC size, whereas for *Fmr1^−/y^* L2/3 PNs responses were unchanged (Supplementary Fig. 2).

To investigate if hyperexcitability is attributable to single L4 to L2/3 PN synaptic responses, we performed a minimal stimulation protocol and found a significant increase in the size of minimally evoked EPSC from *Fmr1^−/y^*compared to *Fmr1^+/y^* L2/3 PNs (*P*=0.007, Fig. 2 D). In the presence of bicuculline the genotypic effect was eliminated (*P*=0.6, Fig. 2 E). To test for changes in L4 presynaptic release properties we performed a paired pulse experiment across four interstimulus intervals (10, 20, 50 & 100 ms) and did not see any effect (ANOVA *P*=1.0, Fig. 2F).

Altogether, this shows that *Fmr1^−/y^* single L4 to L2/3 PN synapses are hyperexcitable compared to controls and this is may be attributable to a disinhibition of *Fmr1^−/y^* L4 to L2/3 PN post synapses (Supplementary Fig. 1).

### Postsynaptic & intrinsic properties of L2/3 PNs are unchanged

Next, we tested for changes in L2/3 PN spontaneous EPSCs (sEPSCs) between the genotypes and found no change in amplitude (*P*=0.8, Fig. 3A). There was a slight significant increase in sEPSC frequency of *Fmr1^−/y^* compared to control L2/3 PNs (*P*=0.047, Fig. 3A).

**Figure 3.**
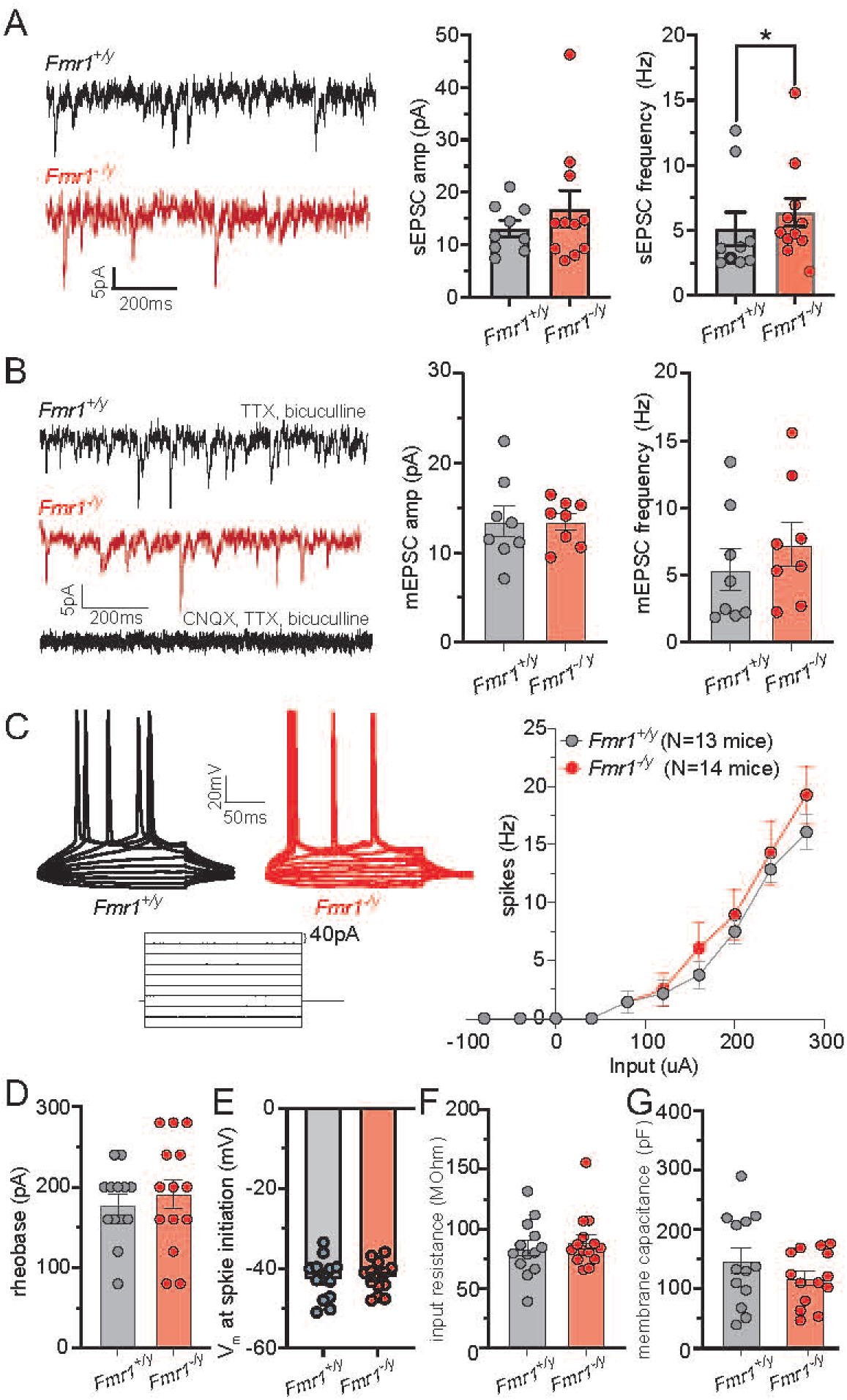
L2/3 PNs intrinsic properties are unchanged in *Fmr1^−/y^* mice. **(A)** Left, representative whole-cell patch-clamp (−70 mV) recordings of sEPSCs for *Fmr1^+/y^* (black) and *Fmr1^−/y^* (red) L2/3 PNs. Middle, mean sEPSC amplitude for *Fmr1^+/y^*(N=9 mice, 13.14+/− 1.52) & *Fmr1^−/y^* (N=11 mice, 16.91 +/− 3.44, *P*=0.8) L2/3 PNs. Right, mean sEPSC frequency for *Fmr1^+/y^*(N=9 mice, 5.10+/− 1.30) & *Fmr1^−/y^* (N=11 mice, 6.39 +/− 1.10, *P*=0.047). **(B)** Left, Representative whole-cell patch-clamp (−70 mV) recordings of *Fmr1^+/y^*(black) and *Fmr1^−/y^* (red) mEPSCs in L2/3 PNs. In the presence of 1μM tetrodotoxin (TTX) and 10μM bicuculline to block GABAA receptors. Middle, mean mEPSC amplitude for *Fmr1^+/y^* (N=9 mice, 13.48+/− 1.69) & *Fmr1^−/y^*(N=11 mice, 13.46 +/− 0.89, p=1.0) L2/3 PNs. Right, mean mEPSC frequency for *Fmr1^+/y^* (N=8 mice, 5.41+/− 1.54) & *Fmr1^−/y^*(N=8 mice, 7.28 +/− 1.64, *P*=0.4) L2/3 PNs. **(C)** Left, representative firing patterns upon current injection steps (40 pA) for *Fmr1^+/y^* & *Fmr1^−/y^*L2/3 PNs. Right, current input (pA) versus mean action potential (AP) frequency (Hz) for *Fmr1^+/y^* (N=13 mice) and *Fmr1^−/y^*(N=14 mice, Two-way ANOVA, time x column p=0.9) L2/3 PNs. **(D)** Mean rheobase for *Fmr1^+/y^* (N=13 mice, 178.5 +/− 12.5) & *Fmr1^−/y^*(N=14 mice, 191.4+/− 18.36, *P*=0.4) L2/3 PNs. **(E)** Mean V_m_ at spike initiation for *Fmr1^+/y^* (N=13 mice, −42.59+/− 1.64) & *Fmr1^−/y^* (N=14, −42.17+/− 1.02, *P*=0.8) L2/3 PNs. **(F)** Mean input resistance for *Fmr1^+/y^* (N=13 mice, 83.77+/− 6.51) & *Fmr1^−/y^* (N=14 mice, 89.15 +/− 6.08, *P*=0.6) L2/3 PNs. **(G)** Mean membrane capacitance for *Fmr1^+/y^* (N=13 mice, 83.77+/− 6.51) & *Fmr1^−/y^* (N=14 mice, 89.15 +/− 6.08, *P*=0.6) L2/3 PNs.

An increase in the size of L4 inputs and impaired plasticity in *Fmr1^−/y^*PNs could be the result of changes in post synaptic glutamate receptor abundance. To test for changes in numbers of postsynaptic glutamate receptors we measured miniature EPSCs (mEPSCs, holding potential - 70 mV) from L2/3 PNs in in the presence of 1μM tetrodotoxin (TTX) and 10 μM bicuculline (Fig. 3B). mEPSCs were eliminated in the presence of 5 μM CNQX, demonstrating that they were due to alpha-amino-3-hydroxy-5-methyl-4-isoxazole propionic acid receptor (AMPAR) activity. We did not find changes in mEPSC amplitude (*P*=1.0) or frequency (*P*=0.4) between the two genotypes (Fig. 3B).

As *Fmr1^−/y^* L4 to L2/3 PN input strength is increased, we examined if this impacted intrinsic cellular excitability. We measured spiking from 40 pA current injection steps and found no differences in spiking frequency, rheobase, membrane potential (V_m_) at spike initiation, input resistance, or membrane capacitance between *Fmr1^−/y^*and control L2/3 PNs (Fig. 3C-D). Altogether, we found no change in postsynaptic AMPA receptor abundance or intrinsic excitability.

### Rhythmic whisker-evoked plasticity is impaired in *Fmr1^−/y^* mice

Single whiskers rhythmically stimulated at the same frequency (8Hz) used in the L4-RES protocol *ex vivo*, previously termed Rhythmic Whisker Stimulation (RWS), in anesthetized mice and awake mice has been shown to induce plasticity in L2/3 neurons ^11,12,19,25^. To monitor whisker-evoked activity of L2/3 neurons in the barrel cortex of S1, we expressed the genetically encoded Ca^2+^ sensor GCaMP6s using adeno-associated viral vectors (AAV1-hsyn-mRuby2-GSG-P2A-GCaMP6s). Single cell Ca^2+^ signals were recorded using two-photon laser scanning microscopy (2P) through a chronically implanted cranial window (Fig. 4A,B). Extracted Ca^2+^ signals were classified as whisker-evoked events when time locked to the whisker stimulus. The stimulation response window was determined by acquisition frame rates and typical GCaMP6s response kinetics ^66^. For each neuron, we calculated the Whisker-evoked response strength (RS; hereafter simply ‘response strength’).

**Figure 4.**
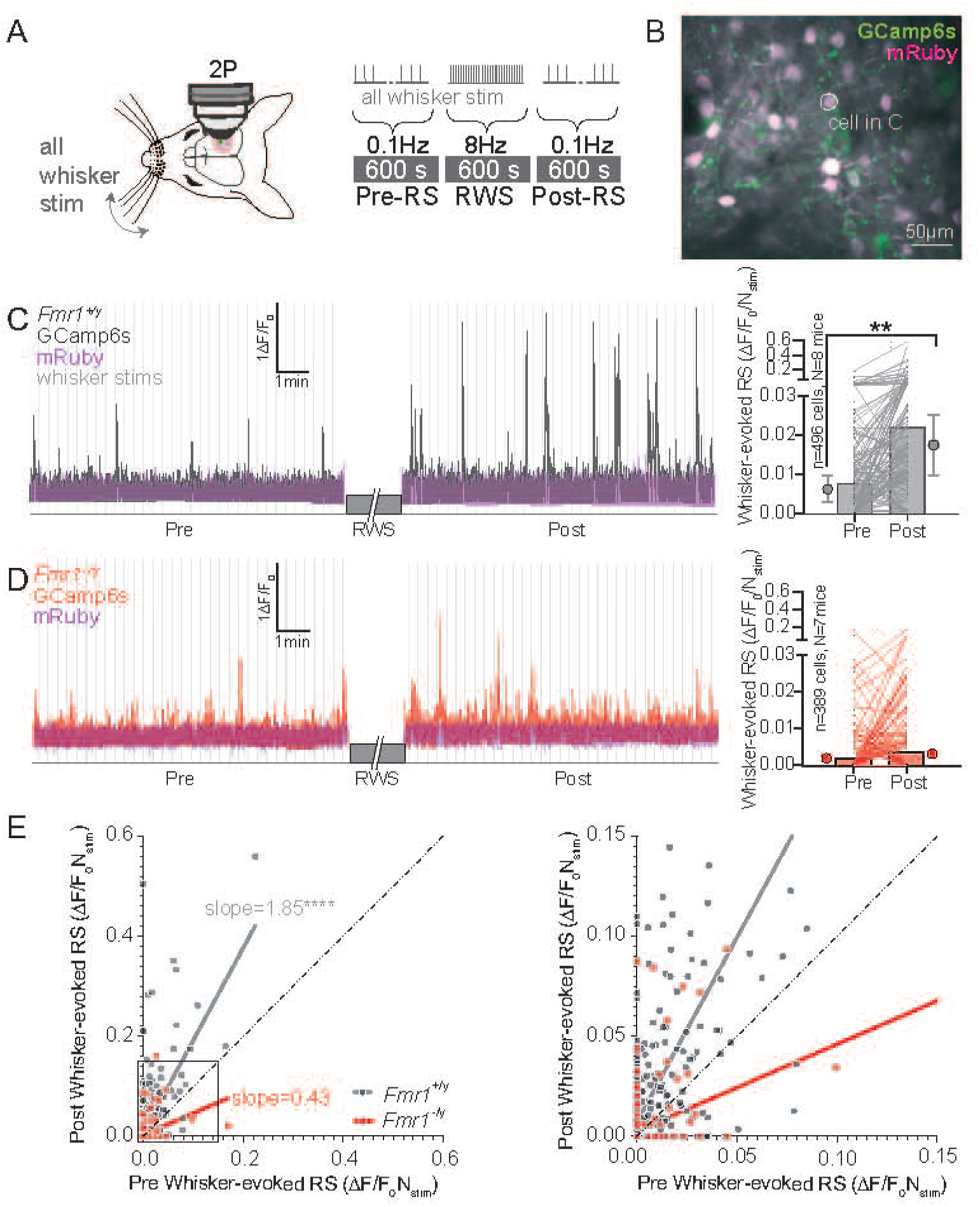
Impaired RWS-evoked plasticity in *Fmr1^−/y^* mice. **(A)** Experimental protocol: all the whiskers stimulated at 0.1 Hz for 10 min (600s) pre- and post-rhythmic whisker stimulation (RWS). RWS is performed at 8Hz for 10 min. while Ca^2+^ imaging is performed using two-photon (2P) microscopy. **(B)** Average 2P image of GCaMP6s-expressing (green) and mRuby (magenta), an activity independent marker of expression, in L2/3 neurons. The signals in **(C)** are from the cell circled. **(C & D)** Left, example trace of the GCamp6s fluorescence, in response to all whisker stimulation (0.1Hz, 10 min) pre- & post RWS. Right, the whisker-evoked response strength (RS, amplitude X whisker-evoked signal probability, (ΔF/F_0_)/Nstim) pre- and post-RWS for *Fmr1^+/y^* (**C**, pre=0.007+/−0.0009, post=0.02+/−0.002, n=496 cells, N=8 mice, *P*=0.008) & *Fmr1^−/y^* (**D**, pre=0.002+/−0.0009, post=0.003+/−0.0009, n=389 cells, N=7 mice, *P*=0.4) L2/3 neurons. **(E)** Left, pre- vs post-RWS RS (ΔF/F_0_)/Nstim) for all cells for *Fmr1^+/y^* (grey, simple linear regression slope=1.9, n=496 cells) and *Fmr1^−/y^* (red, slope=0.4, n=389 cells) mice (significantly non-zero both? P<0.0001. Are slopes equal? *P*<0.001). Comparing slopes (*Fmr1^+/y^* =1.85+/− 0.1, *Fmr1^−/y^* = 0.43+/−0.06, *P*<0.0001). Right, square in left.

We found that RWS of all whiskers under light anaesthesia, medetomidine and midazolam, significantly increased the mean response strength of the whole L2/3 neuronal population of control *Fmr1^+/y^* mice (*P*=0.008, LMM *P*=0.02, see Supplementary Data Fig. 1 for LMM statistics, Fig. 4C). For *Fmr1^−/y^* mice the mean response strength remained unchanged (*P*=0.4, LMM *P*=0.2, Fig. 4D). When we plotted the pre versus post-RWS response strength and ran a simple linear regression analysis, we found for *Fmr1^−/y^* mice the slope deviated below the identity line (slope=0.43; Fig. 4E), whereas for *Fmr1^+/y^* mice the slope was significantly above (slope=1.85, *P*<0.0001). This indicates a significant increase in response strength post RWS, which was not present in *Fmr1^−/y^* mice. Altogether, reinforcing our results *ex vivo* that repetitively evoked plasticity at whisking rates is impaired in *Fmr1^−/y^*mice.

### Persistent cell hypoplasticity is impaired *Fmr1^−/y^* mice

As we observed hyperexcitable L4 to L2/3 PN synaptic responses, we wanted to test if this translated into proportionate changes in whisker-evoked L2/3 neuronal responses in S1 (Fig. 5). To test this, we looked at response strength of ‘active’ cells, termed so by having at least one whisker-evoked event during the pre-RWS period (10 min), and did not find a significant difference between genotypes (*P*=0.8, LMM *P*=0.6, Fig. 5B). When we looked at the proportion of active cells across the entire experiment (30 min) we found *Fmr1^−/y^* cells had a decreased probability trend (binomial test p<0.0001) of being active (16%) compared to *Fmr1^+/y^* (39%) cells, but this was not significant for the LMM (*P*=0.1, Fig. 5C).

**Figure 5.**
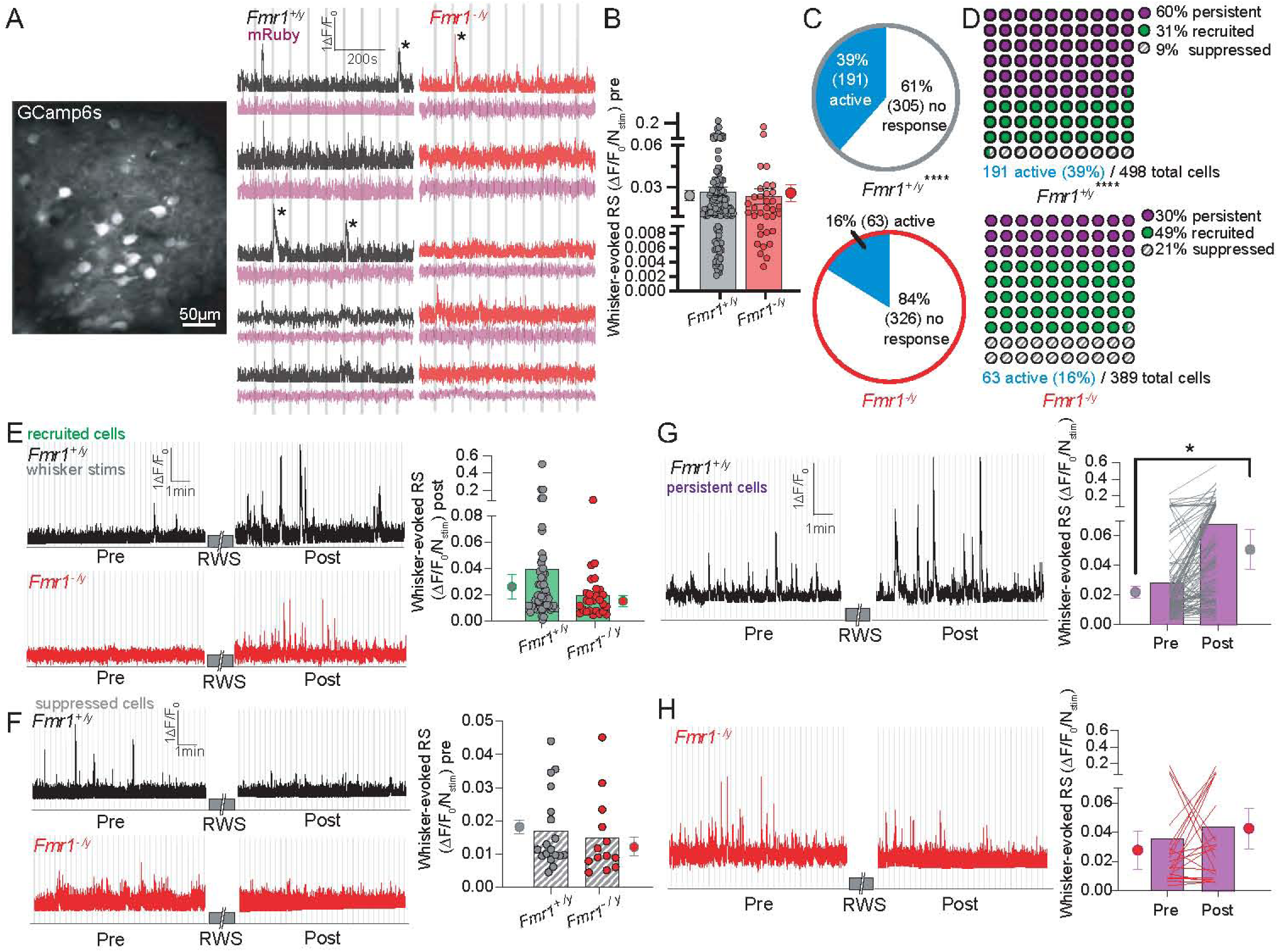
Impaired persistent cell plasticity in *Fmr1^−/y^* mice. **(A)** Left, mean 2P image of GCaMP6s-expressing L2/3 neurons in S1. Right, example traces of the GCamp6s fluorescence in five randomly chosen cells from one *Fmr1^+/y^* and *Fmr1^−/y^* mouse in response to all whisker stimulation (0.1Hz, 10 min). Asterix denotes passing the threshold for a whisker-evoked event. **(B)** Mean whisker-evoked RS for L2/3 neurons active in the pre period in *Fmr1^+/y^*(0.02+/ 0.003, n=138 cells, N=8 mice) & *Fmr1^−/y^* (0.02+/−0.006, n=33 cells, N=5 mice, *P*=0.8) mice. **(C)** Pie charts with the percentage (%) of active and no response cells across the entire RWS protocol for L2/3 neurons from *Fmr1^+/y^* (active: 39%, 191 cells; no response: 61%, 305 cells) and *Fmr^−/y^* (active: 16%, 63 cells; no response: 84%, 326 cells) mice (O vs. E binomial test, *P*<0.0001). **(D)** Parts of whole graphs with the percentages (%) of persistent (purple, 60% & 30%), recruited (green, 31% & 49%), & suppressed (9% & 21%) L2/3 cells for *Fmr1^+/y^* and *Fmr1^−/y^* mice respectively (O vs E Chi-square=79.86, *P*<0.0001). **(E&F)** Left, top *Fmr1^+/y^* & bottom *Fmr1^−/y^*example trace of GCaMP6s fluorescence from L2/3 neurons of which responses were recruited (E) or suppressed (F) post-RWS. Right, the mean whisker-evoked response strength (RS) of recruited L2/3 neurons post-RWS (E) for *Fmr1^+/y^*(0.03+/−0.01, n=59 cells, N=8 mice) & *Fmr1^−/y^* (0.02+/−0.004, n=31 cells, N=11 mice, p=0.2) mice, or suppressed L2/3 neurons (G) pre-RWS for *Fmr1^+/y^*(0.02+/−0.002, n=18 cells, N=8 mice) & *Fmr1^−/y^* (0.01+/−0.003, n=13 cells, N=5 mice, *P*=0.1) mice. **(G, H)** Left, top *Fmr1^+/y^* & bottom *Fmr1^−/y^*example trace of GCaMP6s fluorescence from a L2/3 neuron showing persistent whisker-evoked responses (0.1 Hz, 10 min) pre- and post-RWS. **(G)** Right, bar and pairwise representation for *Fmr1^+/y^* L2/3 neurons (n=114 cells, N=8, p=0.035) pre- (0.02+/−0.004) & post-RWS (0.05+/−0.01) for (0.03+/−0.01), or (H) for *Fmr1^−/y^* (n=19 cells, N=5, *P*=0.3) pre- (0.03+/−0.01) and post-RWS (0.04+/−0.01).

Cortical neuronal populations show a remarkable heterogeneity in sensory-evoked activity ^3,19,67–70^. As we previously observed in awake mice, we found that nearly half of the imaged neuronal population showed one or more PW-evoked Ca^2+^ signals pre- and post-RWS and were termed ‘persistent’ cells (Fig. 5G,H) ^19^. The remaining neurons lacked PW-evoked Ca^2+^ signals pre- and/ or post-RWS. This group was subdivided into those that lacked signals pre-RWS but displayed them at any time point post-RWS – termed ‘recruited’ cells (Fig. 5E), and those that displayed PW-evoked signals pre-RWS but not at any time point post-RWS – termed ‘suppressed’ cells (Fig. 5F). We found that the proportion of persistent (*Fmr1^+/y^*=60% vs *Fmr1^−/y^*=30%), recruited (*Fmr1^+/y^*=31% vs *Fmr1^−/y^*=49%), and suppressed cells (*Fmr1^+/y^*=9% vs *Fmr1^−/y^*=21%) showed a trend of difference between *Fmr1^+/y^* and *Fmr1^−/y^*mice but this was not significant for the GLMM (Chi square *P*<0.0001, Fig. 5 D, Supplemental Data Fig.1). For the recruited subpopulation, the mean post-RWS response strength was similar for both genotypes (*P*=0.2, LMM *P*=0.5, Fig. 5E). For the suppressed subpopulation the mean pre-RWS response strength was also unchanged (*P*=0.1, LMM=0.2, Fig. 5F). Consistent with our overall findings the persistent subpopulation for control mice exhibited a significant increase in response strength upon RWS (*P*=0.035, LMM *P*=8.7e-9, Fig. 5G). In contrast, we observed no change in response strength for *Fmr1^−/y^* mice (*P*=0.3, LMM *P*=0.4, Fig. 5H).

Together, these data show no change in the size or probability whisker-evoked responses between the groups. RWS, however, fails to potentiate whisker-evoked activity in the persistent cell population in L2/3 of *Fmr1^−/y^*mice compared to controls.

### AIS length is decreased in L2/3 of *Fmr1^−/y^* mice

We next wanted to investigate the paradoxical effect of subthreshold hyperexcitability and unchanged or decreased sensory-evoked neuronal activation in L2/3. The AIS is a specialised microdomain, characterized by enrichment of ion channels and specialized cytoskeletal proteins such as Ankyrin-G, where action potential initiation may be proportional to AIS length ^49,53,71^. Previous literature has reported changes in AIS length in rodent models of neurodevelopmental and neurological disorders, including *Fmr1^−/y^* mice ^50,51,54^. To measure changes in AIS length we performed immunohistochemical labelling in L2/3 and L4 of S1 for anti-Ankyrin-G (Neuromab), as an AIS marker, and NeuN (Millipore EMD), a general neuronal marker (Fig. 6A). Z-stack images were acquired using confocal microscopy, and mean length of L2/3 and L4 AISs were measured using ImageJ FIJI. We found for L2/3 AIS lengths were significantly shorter for *Fmr1^−/y^* compared to littermates (*P*=0.001, Fig. 6B,D), whereas AIS lengths in L4 were unchanged (*P*=0.07, Fig. 6A,C,D). We also found the relative frequency distribution of AIS length significantly shifted towards smaller AIS in L2/3 of *Fmr1^−/y^* compared to controls (*P*=0.0006), but not for L4 (*P*=0.9, Fig. 6D). Together, this data may reveal that *Fmr1^−/y^* mice exhibit decreased AIS length in L2/3 but not in L4 compared to controls and this may explain the combined effect of subthreshold hyperexcitability with relatively unchanged or modestly decreased action potential generation.

**Figure 6.**
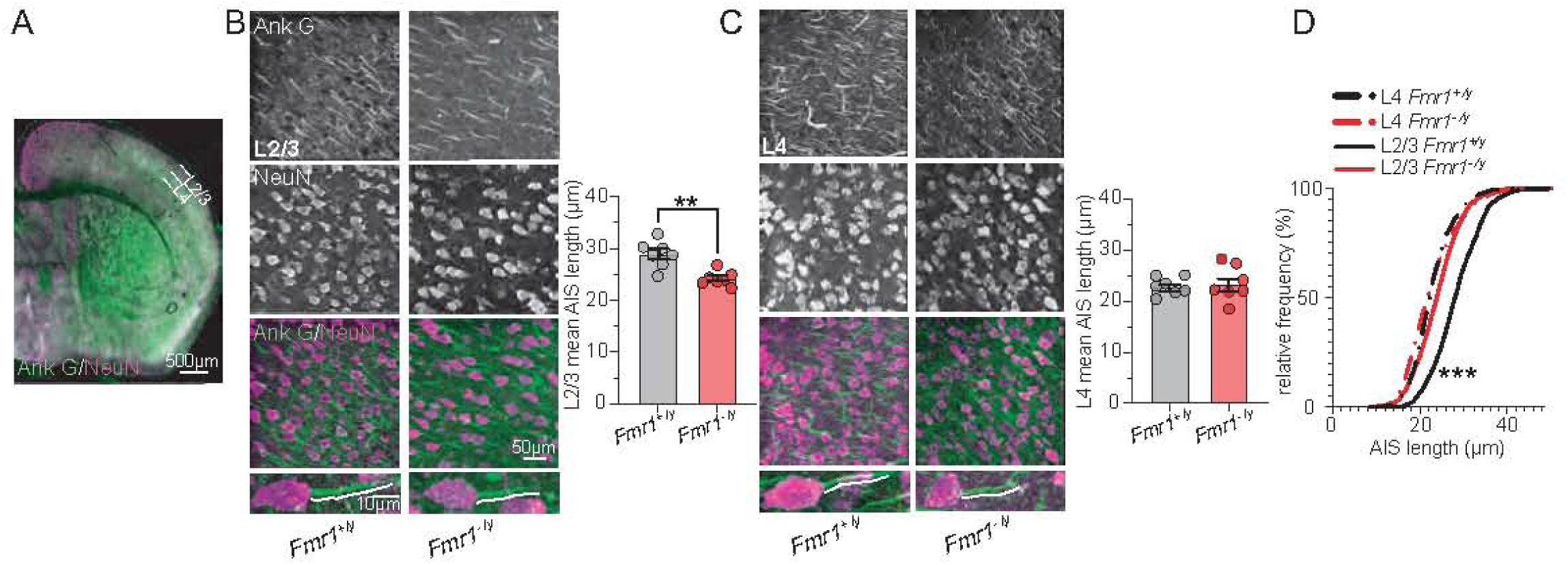
AIS length in L2/3 is decreased in *Fmr1^−/y^* mice. **(A)** Ankryin-G (green) & Anti-NeuN (pink) in an S1 brain slice with analysis in L2/3 & 4 S1. **(B & C)** Left, representative *Fmr1^+/y^* (left) & *Fmr1^−/y^* (right) staining in L2/3 **(B)** or L4 **(C)** of S1 in fixed tissue for Anti-Ankryin-G (top), NeuN (second), overlap (third, white), & magnified (bottom). White dotted line is a representative length measurement for a single Axon Initial Segment (AIS). Right, bar graph of mean AIS length (μm) for L2/3 (*Fmr1^+/y^*=28.9±1, N=7, *Fmr1^−/y^*=24.2±0.5, N=7, p=0.001) or L4 (*Fmr1^+/y^*=22.9±0.6, N=7, *Fmr1^−/y^*=23.5±1.3, N=7, *P*=0.7). **(D)** AIS length versus relative frequency (%) for AIS lengths of *Fmr1^+/y^* and *Fmr1^−/y^* fixed brain slices in L2/3 (KS test, *P*=0.0006) or L4 (*P*=0.9).

### Adaptation to the plasticity protocol is increased in *Fmr1^−/y^* mice

It has previously been shown that the prolonged activation of L2/3 PNs to repetitive stimulation drives a sustained NMDAR-dependent depolarization, which is necessary for LTP expression ^19,25,27^. To determine if loss of FMRP impacts activity during the RES protocol we looked at evoked L4-ES PSP amplitudes from L2/3 PNs during L4-RES stimulation (Fig. 7) ^19,25,27^. We found a significant shortening of the duration (<5 sec) of L2/3 PN activation during L4-RES for *Fmr1^−/y^*mice consistent with an increase in adaptation to sensory stimulation. Whereas for control *Fmr1^+/y^* mice amplitudes remained relatively sustained for the first 5-15 sec (*P*=0.02, Fig. 7A, B, D). Moreover, L2/3 PN cumulative amplitudes were decreased for *Fmr1^−/y^* mice compared to controls (*P*<0.0001; Fig. 7A-C).

**Figure 7.**
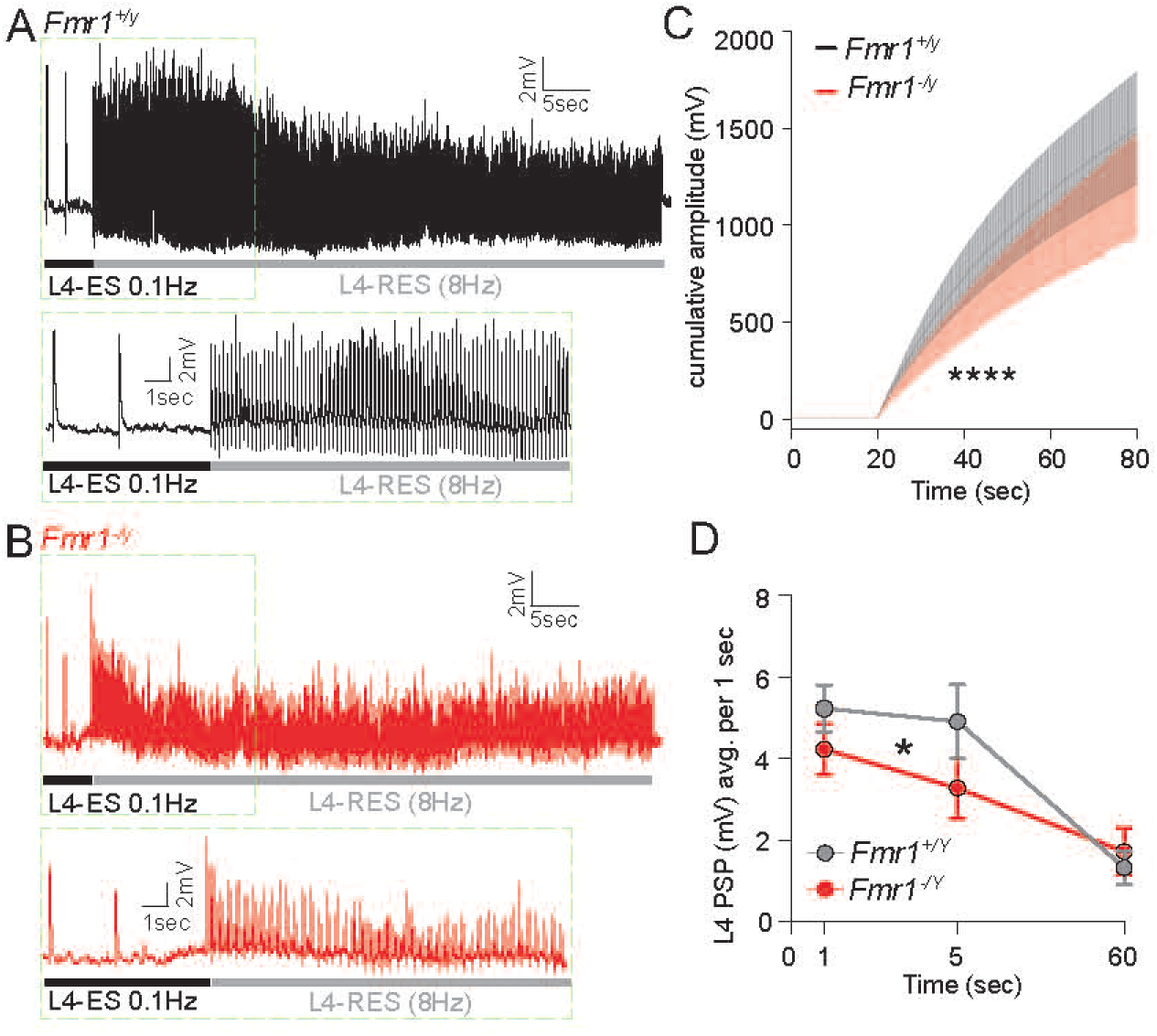
Adaptation to the plasticity protocol is increased in *Fmr1^−/y^* mice. **(A, B)** Representative traces of L4-Rhythmic Electrical Stimulation (L4-RES) for *Fmr1^+/y^* **(A)** & *Fmr1^−/y^* **(B)** L2/3 PNs. Bottom, magnified representative traces for top green dotted box in top of L4-RES *Fmr1^+/y^***(A)** and *Fmr1^−/y^* **(B)** for L2/3 PNs. **(C)** Cumulative L4-ES PSP amplitudes during RPS comparing *Fmr1^+/y^* (N=8 mice) & *Fmr1^−/y^* (N=9 mice, KS test *P*<0.0001). **(D)** Mean L4-ES PSP amplitude across time points (1, 5, 60 sec) during RPS comparing *Fmr1^+/y^* (N=8 mice) & *Fmr1^−/y^* (N=9 mice, Two-way ANOVA, time x genotype, *P*=0.02) L2/3 PNs.

Altogether, mechanistically this may imply that lack of sufficient depolarization, possibly driven by increased adaptation to repetitive sensory stimulation, may underpin hypoplasticity of L2/3 PNs in Fmr1−/y mice.

## Discussion

Repetitive whisker stimulation in rodents under anaesthesia elicits LTP of cortical excitatory synapses ^11,12,19,25^. Here, we repetitively stimulated L4 incoming sensory circuits while recording in whole-cell configuration L2/3 PNs in brain slices from *Fmr1^−/y^* mice and found impaired LTP. Similarly, plasticity was impaired *in vivo* in *Fmr1^−/y^* mice upon a bout of repetitive whisker stimulation as measured by somatic Ca^2+^ signals ^19,25^. Impaired plasticity was largely attributable to a hypoplasticity of the persistent cell population in *Fmr1^−/y^* mice, or cells that are responsive to whisker stimulation across the experiment. Altogether, this strongly suggests that cortical plasticity evoked at physiological whisking frequencies is impaired in *Fmr1^−/y^* mice.

Synaptic L4 to L2/3 PN strength (EPSCs & PSPs) was significantly increased at the level of single synaptic responses. Moreover, we found that significantly less L4-electrical stim current evoked similar PSP sizes compared to littermates. For comparison, L4 to L2/3 PN EPSC size in *Fmr1^−/y^* juveniles (postnatal (P) day 17-23) has been shown to be diminished, this contrast may reflect an emergent developmental phenotype ^72^.

Blocking inhibition in slices from littermate *Fmr1^+/y^*mice increased L4 to L2/3 PN EPSC size and eliminated genotypic differences. Whereas for *Fmr1^−/y^* L2/3 PNs we found no effect, which suggests that there may a loss of inhibitory tone on L4 to L2/3 PN circuit consistent with decreased inhibition. This data builds upon a large body of work demonstrating impaired and altered timing of feedforward inhibition in FXS individuals and *Fmr1^−/y^* rodents ^72–74^.

L4 to L2/3 PN hyperexcitability did not precipitate increased intrinsic spiking or whisker-evoked response strength. This recapitulates previous 2-photon results in juvenile (P14-16) *Fmr1^−/y^* mice compared to controls, which showed no genotypic difference in whisker-evoked activity with significantly fewer *Fmr1^−/y^*neurons responsive to whisker stimulation ^72^. Single unit recordings in similarly aged *Fmr1^−/y^* adults and juveniles (P17-23 mice) have shown decreased firing in response to whisker stimulation, which is not incompatible with our results considering our methods^75,76^

Homeostatic plasticity serves to maintain the neuronal set point optimal for information processing ^77,78^. Our observations suggest that increased synaptic strength does not necessarily increase spiking in *Fmr1^−/y^* mice. Here we find a decrease in L2/3 AIS length in *Fmr1^−/y^* mice, which may account for no change in spiking. Regulation of AIS length to acute changes in activity has recently been challenged, however, altered AIS length has been reported in several models of neurodevelopmental and pathological conditions^50,54,79–81^.

Previously, stimulation of incoming whisker circuits at physiological frequencies and RWS has been shown to evoke in L2/3 PNs a period of sustained NMDAR-mediated depolarization, which is required to elicit LTP ^25,82–85^. Here we found that during the plasticity inducing protocol there is an absence of sustained depolarization in *Fmr1^−/y^* L2/3 PNs compared to controls ^27^. This could be the result of altered sensory adaptation, and this is supported by observations in juvenile (P14-P16 mice) *Fmr1^−/Y^* L2/3 neurons ^75,86^. This has also been observed in similar experiments *in* and *ex vivo* when the POm-higher order thalamus is inhibited or in the presence of NMDAR antagonists, which suggests a similar mechanistic underpinning^25,27^. Future studies examining the relationship between FMRP, dendritic integration, and sensory adaptation mechanisms are required to uncover the mechanism.

The cerebral cortex integrates sensory information with emotional states and internal representations to produce coherent precepts, form associations and execute goal-directed behaviour. Impaired LTP in the sensory cortex suggests that the receptive field properties of cortical PNs may be less flexible and dynamic. This may manifest as measurable changes in tactile behavioural responses in *Fmr1^/−y^* mice and may contribute to altered tactile reactivity in FXS, which should be explored in future ^87^. Altered sensory reactivity is common across many neurological conditions, including neurodevelopmental disorders, intellectual disability, and autism; future studies should determine whether impaired or imbalanced sensory-driven LTP expression in the sensory neocortex constitutes an underlying convergent phenotype^88^.

## Data availability

All datasets will be made available on Edinburgh Datashare. Large primary datasets can be made available upon request.

## Acknowledgements

We thank Nathalie Rochefort, Danai Katsenevaki, and Darren Walsh for their help with software. We thank Nathalie Rochefort for her comments on the manuscript. We thank the Bioresearch and Veterinary Services (BVS) at the University of Edinburgh for the care and maintenance of all animals used in this study.

## Abbreviations

No abbreviations list. Abbreviations should be defined at first use (except for accepted gene/protein symbols). Abbreviations in figures and tables should be defined in the legend.

## Supplementary material

Supplementary material is available at *Brain* online.

## References

1. Feldman DE. Synaptic mechanisms for plasticity in neocortex. Annu Rev Neurosci. 2009;32:33–55. doi:10.1146/annurev.neuro.051508.135516

2. Glazewski S, Chen CM, Silva A, Fox K. Requirement for alpha-CaMKII in experience-dependent plasticity of the barrel cortex. Science. Apr 19 1996;272(5260):421–3. doi:10.1126/science.272.5260.421

3. Margolis DJ, Lütcke H, Schulz K, et al. Reorganization of cortical population activity imaged throughout long-term sensory deprivation. Nat Neurosci. Nov 2012;15(11):1539–46. doi:10.1038/nn.3240

4. Hardingham N, Wright N, Dachtler J, Fox K. Sensory deprivation unmasks a PKA-dependent synaptic plasticity mechanism that operates in parallel with CaMKII. Neuron. Dec 10 2008;60(5):861–74. doi:10.1016/j.neuron.2008.10.018

5. Jacob V, Brasier DJ, Erchova I, Feldman D, Shulz DE. Spike timing-dependent synaptic depression in the in vivo barrel cortex of the rat. J Neurosci. Feb 7 2007;27(6):1271–84. doi:10.1523/jneurosci.4264-06.2007

6. Pawlak V, Greenberg DS, Sprekeler H, Gerstner W, Kerr JN. Changing the responses of cortical neurons from sub- to suprathreshold using single spikes in vivo. Elife. Jan 22 2013;2:e00012. doi:10.7554/eLife.00012

7. El-Boustani S, Ip JPK, Breton-Provencher V, et al. Locally coordinated synaptic plasticity of visual cortex neurons in vivo. Science. Jun 22 2018;360(6395):1349–1354. doi:10.1126/science.aao0862

8. Gambino F, Holtmaat A. Spike-timing-dependent potentiation of sensory surround in the somatosensory cortex is facilitated by deprivation-mediated disinhibition. Neuron. Aug 9 2012;75(3):490–502. doi:10.1016/j.neuron.2012.05.020

9. Clapp WC, Kirk IJ, Hamm JP, Shepherd D, Teyler TJ. Induction of LTP in the human auditory cortex by sensory stimulation. Eur J Neurosci. Sep 2005;22(5):1135–40. doi:10.1111/j.1460-9568.2005.04293.x

10. Frenkel MY, Sawtell NB, Diogo AC, Yoon B, Neve RL, Bear MF. Instructive effect of visual experience in mouse visual cortex. Neuron. Aug 3 2006;51(3):339–49. doi:10.1016/j.neuron.2006.06.026

11. Mégevand P, Troncoso E, Quairiaux C, Muller D, Michel CM, Kiss JZ. Long-Term Plasticity in Mouse Sensorimotor Circuits after Rhythmic Whisker Stimulation. The Journal of Neuroscience. 2009;29(16):5326–5335. doi:10.1523/jneurosci.5965-08.2009

12. Han Y, Huang MD, Sun ML, Duan S, Yu YQ. Long-Term Synaptic Plasticity in Rat Barrel Cortex. Cereb Cortex. Sep 2015;25(9):2741–51. doi:10.1093/cercor/bhu071

13. Sanders PJ, Thompson B, Corballis PM, Maslin M, Searchfield GD. A review of plasticity induced by auditory and visual tetanic stimulation in humans. Eur J Neurosci. Aug 2018;48(4):2084–2097. doi:10.1111/ejn.14080

14. Lengali L, Hippe J, Hatlestad-Hall C, Rygvold TW, Sneve MH, Andersson S. Sensory-Induced Human LTP-Like Synaptic Plasticity - Using Visual Evoked Potentials to Explore the Relation Between LTP-Like Synaptic Plasticity and Visual Perceptual Learning. Front Hum Neurosci. 2021;15:684573. doi:10.3389/fnhum.2021.684573

15. Marzoll A, Shibata K, Toyoizumi T, Chavva I, Watanabe T. Decrease in signal-related activity by visual training and repetitive visual stimulation. iScience. Dec 22 2022;25(12):105492. doi:10.1016/j.isci.2022.105492

16. Kato HK, Gillet SN, Isaacson JS. Flexible Sensory Representations in Auditory Cortex Driven by Behavioral Relevance. Neuron. Dec 2 2015;88(5):1027–1039. doi:10.1016/j.neuron.2015.10.024

17. Chéreau R, Bawa T, Fodoulian L, Carleton A, Pagès S, Holtmaat A. Dynamic perceptual feature selectivity in primary somatosensory cortex upon reversal learning. Nat Commun. Jun 26 2020;11(1):3245. doi:10.1038/s41467-020-17005-x

18. Kowalewski R, Kattenstroth JC, Kalisch T, Dinse HR. Improved acuity and dexterity but unchanged touch and pain thresholds following repetitive sensory stimulation of the fingers. Neural Plast. 2012;2012:974504. doi:10.1155/2012/974504

19. Williams LE, Küffer L, Bawa T, Husi E, Pagès S, Holtmaat A. Repetitive Sensory Stimulation Potentiates and Recruits Sensory-Evoked Cortical Population Activity. The Journal of Neuroscience. 2025;45(4):e2189232024. doi:10.1523/jneurosci.2189-23.2024

20. Bliss TV, Collingridge GL. A synaptic model of memory: long-term potentiation in the hippocampus. Nature. Jan 7 1993;361(6407):31–9. doi:10.1038/361031a0

21. Carvell GE, Simons DJ. Biometric analyses of vibrissal tactile discrimination in the rat. J Neurosci. Aug 1990;10(8):2638–48. doi:10.1523/jneurosci.10-08-02638.1990

22. Wolfe J, Hill DN, Pahlavan S, Drew PJ, Kleinfeld D, Feldman DE. Texture coding in the rat whisker system: slip-stick versus differential resonance. PLoS Biol. Aug 26 2008;6(8):e215. doi:10.1371/journal.pbio.0060215

23. Fee MS, Mitra PP, Kleinfeld D. Central versus peripheral determinants of patterned spike activity in rat vibrissa cortex during whisking. J Neurophysiol. Aug 1997;78(2):1144–9. doi:10.1152/jn.1997.78.2.1144

24. Crochet S, Petersen CC. Correlating whisker behavior with membrane potential in barrel cortex of awake mice. Nat Neurosci. May 2006;9(5):608–10. doi:10.1038/nn1690

25. Gambino F, Pagès S, Kehayas V, et al. Sensory-evoked LTP driven by dendritic plateau potentials in vivo. Nature. Nov 6 2014;515(7525):116–9. doi:10.1038/nature13664

26. Roth MM, Dahmen JC, Muir DR, Imhof F, Martini FJ, Hofer SB. Thalamic nuclei convey diverse contextual information to layer 1 of visual cortex. Nature Neuroscience. 2016/02/01 2016;19(2):299–307. doi:10.1038/nn.4197

27. Williams LE, Holtmaat A. Higher-Order Thalamocortical Inputs Gate Synaptic Long-Term Potentiation via Disinhibition. Neuron. Jan 2 2019;101(1):91–102.e4. doi:10.1016/j.neuron.2018.10.049

28. Suzuki M, Larkum ME. General Anesthesia Decouples Cortical Pyramidal Neurons. Cell. Feb 20 2020;180(4):666–676.e13. doi:10.1016/j.cell.2020.01.024

29. Urbain N, Salin Paul A, Libourel P-A, Comte J-C, Gentet Luc J, Petersen Carl CH. Whisking-Related Changes in Neuronal Firing and Membrane Potential Dynamics in the Somatosensory Thalamus of Awake Mice. Cell Reports. 2015/10/27/ 2015;13(4):647–656. 10.1016/j.celrep.2015.09.029

30. Nakajima M, Halassa MM. Thalamic control of functional cortical connectivity. Current Opinion in Neurobiology. 2017/06/01/ 2017;44:127–131. 10.1016/j.conb.2017.04.001

31. Chéreau R, Williams LE, Bawa T, Holtmaat A. Circuit mechanisms for cortical plasticity and learning. Semin Cell Dev Biol. May 2022;125:68–75. doi:10.1016/j.semcdb.2021.07.012

32. Hagerman RJ, Staley LW, O’Conner R, et al. Learning-disabled Males With a Fragile X CGG Expansion in the Upper Premutation Size Range. Pediatrics. 1996;97(1):122–126. doi:10.1542/peds.97.1.122

33. Salcedo-Arellano MJ, Dufour B, McLennan Y, Martinez-Cerdeno V, Hagerman R. Fragile X syndrome and associated disorders: Clinical aspects and pathology. Neurobiology of Disease. Mar 2020;136 doi:ARTN 104740 10.1016/j.nbd.2020.104740

34. Crawford DC, Acuña JM, Sherman SL. FMR1 and the fragile X syndrome: human genome epidemiology review. Genet Med. Sep-Oct 2001;3(5):359–71. doi:10.1097/00125817-200109000-00006

35. Costa-Mattioli M, Sossin WS, Klann E, Sonenberg N. Translational Control of Long-Lasting Synaptic Plasticity and Memory. Neuron. Jan 15 2009;61(1):10–26. doi:10.1016/j.neuron.2008.10.055

36. Bassell GJ, Warren ST. Fragile X syndrome: loss of local mRNA regulation alters synaptic development and function. Neuron. Oct 23 2008;60(2):201–14. doi:10.1016/j.neuron.2008.10.004

37. Darnell JC, Van Driesche SJ, Zhang CL, et al. FMRP Stalls Ribosomal Translocation on mRNAs Linked to Synaptic Function and Autism. Cell. Jul 22 2011;146(2):247–261. doi:10.1016/j.cell.2011.06.013

38. Zalfa F, Giorgi M, Primerano B, et al. The fragile X syndrome protein FMRP associates with RNA and regulates the translation of specific mRNAs at Synapses. Cell. Feb 7 2003;112(3):317–327. doi:Doi 10.1016/S0092-8674(03)00079-5

39. Robertson CE, Baron-Cohen S. Sensory perception in autism. Nat Rev Neurosci. Nov 2017;18(11):671–684. doi:10.1038/nrn.2017.112

40. Marco EJ, Hinkley LB, Hill SS, Nagarajan SS. Sensory processing in autism: a review of neurophysiologic findings. Pediatr Res. May 2011;69(5 Pt 2):48r–54r. doi:10.1203/PDR.0b013e3182130c54

41. Crane L, Goddard L, Pring L. Sensory processing in adults with autism spectrum disorders. Autism. 2009/05/01 2009;13(3):215–228. doi:10.1177/1362361309103794

42. Rogers SJ, Hepburn S, Wehner E. Parent reports of sensory symptoms in toddlers with autism and those with other developmental disorders. J Autism Dev Disord. Dec 2003;33(6):631–42 doi:10.1023/b:jadd.0000006000.38991.a7

43. The Dutch-Belgian Fragile XC, Bakker CE, Verheij C, et al. Fmr1 knockout mice: A model to study fragile X mental retardation. Cell. 1994/07/15/ 1994;78(1):23–33. 10.1016/0092-8674(94)90569-X

44. Contractor A, Klyachko Vitaly A, Portera-Cailliau C. Altered Neuronal and Circuit Excitability in Fragile X Syndrome. Neuron. 2015/08/19/ 2015;87(4):699–715. 10.1016/j.neuron.2015.06.017

45. Dölen G, Osterweil E, Rao BSS, et al. Correction of Fragile X Syndrome in Mice. Neuron. 2007/12/20/ 2007;56(6):955–962. 10.1016/j.neuron.2007.12.001

46. Musumeci SA, Bosco P, Calabrese G, et al. Audiogenic Seizures Susceptibility in Transgenic Mice with Fragile X Syndrome. Epilepsia. 2000/12/01 2000;41(1):19–23. 10.1111/j.1528-1157.2000.tb01499.x

47. Osterweil EK, Krueger DD, Reinhold K, Bear MF. Hypersensitivity to mGluR5 and ERK1/2 leads to excessive protein synthesis in the hippocampus of a mouse model of fragile X syndrome. Article. Journal of Neuroscience. 2010;30(46):15616–15627. doi:10.1523/JNEUROSCI.3888-10.2010

48. Bernardet M, Crusio WE. Fmr1 KO mice as a possible model of autistic features. ScientificWorldJournal. Sep 20 2006;6:1164–76. doi:10.1100/tsw.2006.220

49. Kole MH, Stuart GJ. Signal processing in the axon initial segment. Neuron. Jan 26 2012;73(2):235–47. doi:10.1016/j.neuron.2012.01.007

50. Kaphzan H, Buffington SA, Jung JI, Rasband MN, Klann E. Alterations in intrinsic membrane properties and the axon initial segment in a mouse model of Angelman syndrome. J Neurosci. Nov 30 2011;31(48):17637–48. doi:10.1523/jneurosci.4162-11.2011

51. Usui N, Tian X, Harigai W, et al. Length impairments of the axon initial segment in rodent models of attention-deficit hyperactivity disorder and autism spectrum disorder. Neurochemistry International. 2022/02/01/ 2022;153:105273. 10.1016/j.neuint.2021.105273

52. Ogawa Y, Lim BC, George S, et al. Antibody-directed extracellular proximity biotinylation reveals that Contactin-1 regulates axo-axonic innervation of axon initial segments. Nature Communications. 2023/10/26 2023;14(1):6797. doi:10.1038/s41467-023-42273-8

53. Kole MHP, Ilschner SU, Kampa BM, Williams SR, Ruben PC, Stuart GJ. Action potential generation requires a high sodium channel density in the axon initial segment. Nature Neuroscience. 2008/02/01 2008;11(2):178–186. doi:10.1038/nn2040

54. Booker SA, Simões de Oliveira L, Anstey NJ, et al. Input-Output Relationship of CA1 Pyramidal Neurons Reveals Intact Homeostatic Mechanisms in a Mouse Model of Fragile X Syndrome. Cell Rep. Aug 11 2020;32(6):107988. doi:10.1016/j.celrep.2020.107988

55. Ting JT, Daigle TL, Chen Q, Feng G. Acute brain slice methods for adult and aging animals: application of targeted patch clamp analysis and optogenetics. Methods Mol Biol. 2014;1183:221–42. doi:10.1007/978-1-4939-1096-0_14

56. Agmon A, Connors BW. Thalamocortical responses of mouse somatosensory (barrel) cortexin vitro. Neuroscience. 1991/01/01/ 1991;41(2):365–379. 10.1016/0306-4522(91)90333-J

57. Isaac JTR, Oliet SHR, Hjelmstad GO, Nicoll RA, Malenka RC. Expression mechanisms of long-term potentiation in the hippocampus. Journal of Physiology-Paris. 1996/01/01/ 1996;90(5):299–303. 10.1016/S0928-4257(97)87901-6

58. Oliveira LS, Sumera A, Booker SA. Repeated whole-cell patch-clamp recording from CA1 pyramidal cells in rodent hippocampal slices followed by axon initial segment labeling. STAR Protocols. 2021/03/19/ 2021;2(1):100336. 10.1016/j.xpro.2021.100336

59. Holtmaat A, Bonhoeffer T, Chow DK, et al. Long-term, high-resolution imaging in the mouse neocortex through a chronic cranial window. Nat Protoc. 2009;4(8):1128–44. doi:10.1038/nprot.2009.89

60. Pnevmatikakis EA, Giovannucci A. NoRMCorre: An online algorithm for piecewise rigid motion correction of calcium imaging data. J Neurosci Methods. Nov 1 2017;291:83–94. doi:10.1016/j.jneumeth.2017.07.031

61. Keemink SW, Lowe SC, Pakan JMP, Dylda E, van Rossum MCW, Rochefort NL. FISSA: A neuropil decontamination toolbox for calcium imaging signals. Sci Rep. Feb 22 2018;8(1):3493. doi:10.1038/s41598-018-21640-2

62. Williams LE, Featherstone DE. Regulation of hippocampal synaptic strength by glial xCT. J Neurosci. Nov 26 2014;34(48):16093–102. doi:10.1523/jneurosci.1267-14.2014

63. Delacour J, Houcine O, Costa JC. Modifications of the responses of barrel field neurons to vibrissal stimulation during theta in the awake and undrugged rat. Neuroscience. 1990/01/01/ 1990;37(1):237–243. 10.1016/0306-4522(90)90209-M

64. Aroniadou-Anderjaska V, Keller A. LTP in the barrel cortex of adult rats. Neuroreport. Nov 27 1995;6(17):2297–300. doi:10.1097/00001756-199511270-00007

65. Castro-Alamancos MA, Donoghue JP, Connors BW. Different forms of synaptic plasticity in somatosensory and motor areas of the neocortex. The Journal of Neuroscience. 1995;15(7):5324. doi:10.1523/JNEUROSCI.15-07-05324.1995

66. Chen T-W, Wardill TJ, Sun Y, et al. Ultrasensitive fluorescent proteins for imaging neuronal activity. Nature. 2013/07/01 2013;499(7458):295–300. doi:10.1038/nature12354

67. Sato TR, Gray NW, Mainen ZF, Svoboda K. The functional microarchitecture of the mouse barrel cortex. PLoS Biol. Jul 2007;5(7):e189. doi:10.1371/journal.pbio.0050189

68. Brecht M, Roth A, Sakmann B. Dynamic receptive fields of reconstructed pyramidal cells in layers 3 and 2 of rat somatosensory barrel cortex. J Physiol. Nov 15 2003;553(Pt 1):243–65. doi:10.1113/jphysiol.2003.044222

69. Crochet S, Poulet JF, Kremer Y, Petersen CC. Synaptic mechanisms underlying sparse coding of active touch. Neuron. Mar 24 2011;69(6):1160–75. doi:10.1016/j.neuron.2011.02.022

70. Kerr JN, Greenberg D, Helmchen F. Imaging input and output of neocortical networks in vivo. Proc Natl Acad Sci U S A. Sep 27 2005;102(39):14063–8. doi:10.1073/pnas.0506029102

71. Ogawa Y, Rasband MN. The functional organization and assembly of the axon initial segment. Curr Opin Neurobiol. Jun 2008;18(3):307–13. doi:10.1016/j.conb.2008.08.008

72. Antoine MW, Langberg T, Schnepel P, Feldman DE. Increased Excitation-Inhibition Ratio Stabilizes Synapse and Circuit Excitability in Four Autism Mouse Models. Neuron. Feb 20 2019;101(4):648–661.e4. doi:10.1016/j.neuron.2018.12.026

73. Goel A, Cantu DA, Guilfoyle J, et al. Impaired perceptual learning in a mouse model of Fragile X syndrome is mediated by parvalbumin neuron dysfunction and is reversible. Nat Neurosci. Oct 2018;21(10):1404–1411. doi:10.1038/s41593-018-0231-0

74. Domanski APF, Booker SA, Wyllie DJA, Isaac JTR, Kind PC. Cellular and synaptic phenotypes lead to disrupted information processing in Fmr1-KO mouse layer 4 barrel cortex. Nat Commun. Oct 23 2019;10(1):4814. doi:10.1038/s41467-019-12736-y

75. He CX, Cantu DA, Mantri SS, Zeiger WA, Goel A, Portera-Cailliau C. Tactile Defensiveness and Impaired Adaptation of Neuronal Activity in the Fmr1 Knock-Out Mouse Model of Autism. J Neurosci. Jul 5 2017;37(27):6475–6487. doi:10.1523/jneurosci.0651-17.2017

76. Rupprecht P, Carta S, Hoffmann A, et al. A database and deep learning toolbox for noise-optimized, generalized spike inference from calcium imaging. Nature Neuroscience. 2021/09/01 2021;24(9):1324–1337. doi:10.1038/s41593-021-00895-5

77. Marder E, Goaillard J-M. Variability, compensation and homeostasis in neuron and network function. Nature Reviews Neuroscience. 2006/07/01 2006;7(7):563–574. doi:10.1038/nrn1949

78. Chen L, Li X, Tjia M, Thapliyal S. Homeostatic plasticity and excitation-inhibition balance: The good, the bad, and the ugly. Current Opinion in Neurobiology. 2022/08/01/ 2022;75:102553. 10.1016/j.conb.2022.102553

79. Sumera A, Oliveira LS, Kwiatkowska A, et al. Absence of short-term axon initial segment plasticity in human, mouse, and rat cortical circuits. bioRxiv. 2025:2025.09.01.672986. doi:10.1101/2025.09.01.672986

80. Baalman KL, Cotton RJ, Rasband SN, Rasband MN. Blast wave exposure impairs memory and decreases axon initial segment length. J Neurotrauma. May 1 2013;30(9):741–51. doi:10.1089/neu.2012.2478

81. Hamada MS, Kole MH. Myelin loss and axonal ion channel adaptations associated with gray matter neuronal hyperexcitability. J Neurosci. May 6 2015;35(18):7272–86. doi:10.1523/jneurosci.4747-14.2015

82. Lavzin M, Rapoport S, Polsky A, Garion L, Schiller J. Nonlinear dendritic processing determines angular tuning of barrel cortex neurons in vivo. Nature. Oct 18 2012;490(7420):397–401. doi:10.1038/nature11451

83. Palmer LM, Shai AS, Reeve JE, Anderson HL, Paulsen O, Larkum ME. NMDA spikes enhance action potential generation during sensory input. Nat Neurosci. Mar 2014;17(3):383–90. doi:10.1038/nn.3646

84. Pagès S, Chenouard N, Chéreau R, Kouskoff V, Gambino F, Holtmaat A. An increase in dendritic plateau potentials is associated with experience-dependent cortical map reorganization. Proceedings of the National Academy of Sciences. 2021/03/02 2021;118(9):e2024920118. doi:10.1073/pnas.2024920118

85. Xu NL, Harnett MT, Williams SR, et al. Nonlinear dendritic integration of sensory and motor input during an active sensing task. Nature. Dec 13 2012;492(7428):247–51. doi:10.1038/nature11601

86. Whitmire CJ, Stanley GB. Rapid Sensory Adaptation Redux: A Circuit Perspective. Neuron. Oct 19 2016;92(2):298–315. doi:10.1016/j.neuron.2016.09.046

87. Sachidhanandam S, Sreenivasan V, Kyriakatos A, Kremer Y, Petersen CCH. Membrane potential correlates of sensory perception in mouse barrel cortex. Nature Neuroscience. 2013/11/01 2013;16(11):1671–1677. doi:10.1038/nn.3532

88. Cascio CJ. Somatosensory processing in neurodevelopmental disorders. J Neurodev Disord. Jun 2010;2(2):62–9. doi:10.1007/s11689-010-9046-3

