## Supplementary Figures for "Hypoplasticity in sensory-driven necortical circuits of Fragile X mice"

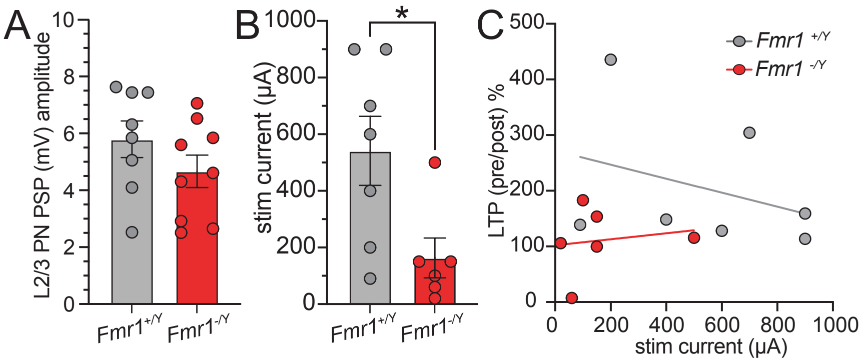


**Supplementary Figure 1. Pre L4-L2/3 PN PSP size between genotypes.**
(A) Mean pre L2/3 PN PSP amplitude bar graph & mouse averages (circles) for *Fmr1*$^{+/Y}$ (N=8 mice, 5.79+/-0.64) & *Fmr1*$^{-/Y}$ (N=9 mice, 4.67+/-0.57, p=0.2).
(B) L4-Electrical Stim (ES) current bar graph & mouse averages for *Fmr1*$^{+/Y}$ (N=7 mice, 541+/-122) & *Fmr1*$^{-/Y}$ (N=6 mice, 163+/-70, p=0.03).
(C) L4-ES stim current versus LTP (pre/post) % linear regression *Fmr1*$^{+/Y}$ (N=7 mice, slope=-0.13, zero slope? p=0.5; Person r, p=0.8, grey) & *Fmr1*$^{-/Y}$ (N=6 mice, slope=0.05, zero slope? p=0.8; Person r, p=0.5, red).

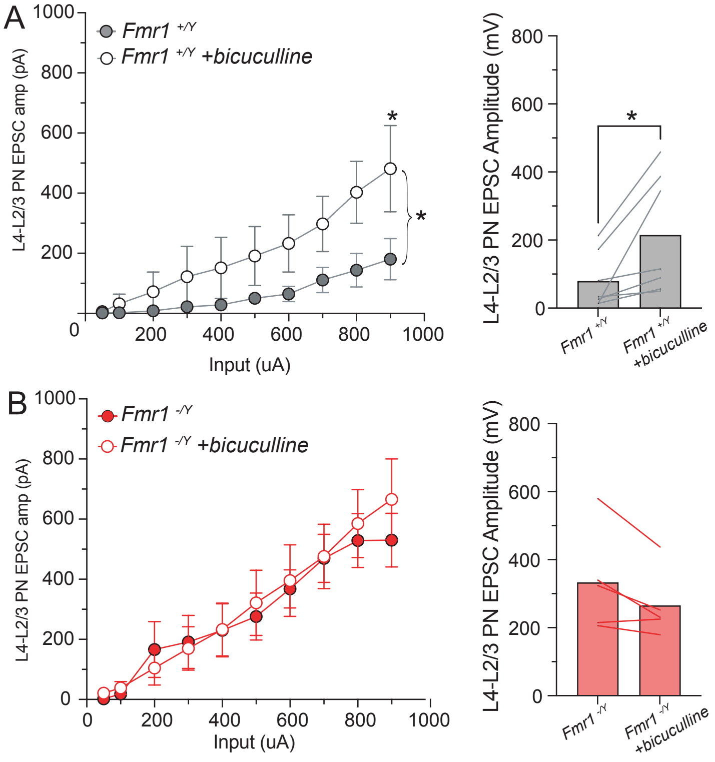


**Supplementary Figure 2. Loss of inhibition does not alter *Fmr1*$^{-/y}$ L4-L2/3 PN EPSC size across various stimulation inputs.**

(A) Left, L4-L2/3 PN EPSCs across various L4-Electrical stimulation (ES) inputs (50, 100-900µA) for *Fmr1*$^{+/y}$ & *Fmr1*$^{+/y}$ +bicuculline (10µM, Two-way ANOVA, time x genotype, p=0.02, for Sidak's mulitple comparisons test 900µA p=0.03). Right, mean L4-L2/3 PN half-max amplitude bar graph and mouse pairwise comparisons for *Fmr1*$^{+/y}$ (N=7 mice, 79.27+/- 30.9) & *Fmr1*$^{+/y}$ +bicuculline (N=7 mice, 214.4 +/- 66.15, p=0.02).

(B) Left, L4-L2/3 PN EPSCs across various L4-ES inputs (50, 100-900µA) for *Fmr1*$^{-/y}$ & *Fmr1*$^{-/y}$ +bicuculline (Two-way ANOVA, time x genotype, p=1.0). Right, mean L4-L2/3 PN half-max amplitude bar graph and mouse pairwise comparisons for *Fmr1*$^{-/y}$ (N=5 mice, 332.9+/- 67.4) & *Fmr1*$^{-/y}$ +bicuculline (N=5 mice, 265.0 +/- 44.7.15, p=0.07).
